# Metabolic consequences of polyphosphate synthesis and imminent phosphate limitation

**DOI:** 10.1101/2022.11.22.517608

**Authors:** Geun-Don Kim, Danye Qiu, Henning Jessen, Andreas Mayer

## Abstract

Cells stabilize intracellular inorganic phosphate (P_i_) to compromise between large biosynthetic needs and detrimental bioenergetic effects of P_i_. P_i_ homeostasis in eukaryotes employs SPXs domains, which are receptors for inositol pyrophosphates. We explored how polymerization and storage of Pi in acidocalcisome-like vacuoles supports S. cerevisiae metabolism and how these cells recognize P_i_ scarcity. Whereas P_i_ starvation affects numerous metabolic pathways, beginning P_i_ scarcity affects few metabolites. These include inositol pyrophosphates and ATP, a low-affinity substrate for inositol pyrophosphate-synthesizing kinases. Declining ATP and inositol pyrophosphates may thus be indicators of impending P_i_ limitation. Actual P_i_ starvation triggers accumulation of the purine synthesis intermediate 5- aminoimidazole-4-carboxamide ribonucleotide (AICAR), which activates P_i_-dependent transcription factors. Cells lacking polyphosphate show P_i_ starvation features already under P_i_-replete conditions, suggesting that vacuolar polyphosphate supplies P_i_ for metabolism even when P_i_ is abundant. However, polyphosphate deficiency also generates unique metabolic changes that are not observed in starving wildtype cells. Polyphosphate in acidocalcisome-like vacuoles may hence be more than a global phosphate reserve and channel P_i_ to preferred cellular processes.

**Abstract importance:** Cells must strike a delicate balance between the high demand of inorganic phosphate (Pi) for synthesizing nucleic acids and phospholipids, and its detrimental bioenergetic effects by reducing the free energy of nucleotide hydrolysis. The latter may stall metabolism. Therefore, microorganisms manage the import and export of phosphate, its conversion into osmotically inactive inorganic polyphosphates, and their storage in dedicated organelles, acidocalcisomes. Here, we provide novel insights into metabolic changes that cells may use to signal declining phosphate availability in the cytosol and differentiate it from actual phosphate starvation. We also analyze the role of acidocalcisome-like organelles in phosphate homeostasis. This uncovers an unexpected role of the polyphosphate pool in these organelles under phosphate-rich conditions, indicating that its metabolic roles go beyond that of a phosphate reserve for surviving starvation.

## Introduction

Inorganic phosphate (P_i_) is an essential nutrient. It is a component of lipids and nucleic acids, controls the activity of proteins through covalent modification, and it serves as an energy transducer when integrated into nucleotides driving many endergonic biochemical reactions. Therefore, perturbed P_i_ homeostasis strongly affects growth and development of various living organisms (1–4). Eukaryotic cells have developed multi-layered systems to and maintain cytosolic P_i_ concentration within a suitable range despite fluctuating environmental conditions. Shortage of intracellular P_i_ was proposed to be signaled through a dedicated signaling pathway for <u>in</u>tracellular <u>pho</u>sphate reception and <u>s</u>ignaling (INPHORS), where the level of cytosolic P_i_ is translated into changes of inositol pyrophosphates (InsPPs), which then bind to SPX (<u>S</u>yg1/<u>P</u>ho81/<u>X</u>pr1) receptor domains (5, 6). These domains form part of or interact with numerous cellular proteins that control transcription or mediate uptake, secretion, storage, and recycling of phosphate, (7, 8). It is expected that INPHORS coordinates these systems such that cytosolic P_i_ concentration is maintained in a viable range.

In S. cerevisiae, INPHORS triggers the PHO pathway, the transcriptional phosphate starvation response that controls expression of genes for P_i_ scavenging, uptake, and recycling (Chabert et al., manuscript in preparation). The PHO pathway in yeast is controlled through the transcription factor Pho4, which accumulates in the nucleus upon P_i_ starvation. It interacts with another transcription factor, Pho2, to activate many P_i_-dependent genes. The interaction between Pho2 and Pho4 is impaired by phosphorylation of Pho4 through Pho85/Pho80 kinase (9, 10). Pho4-Pho2 interaction is stimulated by an intermediate of purine synthesis, AICAR, but it is unknown whether AICAR levels respond to P_i_ availability (11–13).

Many organisms developed strategies to cope with a gradual decline (scarcity) or final absence (starvation) of phosphate, which then becomes growth-limiting (1–4). Budding yeast induces its starvation response in at least two stages depending on the degree of P_i_ depletion (14–16). Phosphate scarcity activates P_i_ scavenging from the environment and maintains growth. Acid phosphatases, such as Pho5, are expressed and secreted to liberate P_i_ from external P_i_-containing molecule (17). As a further measure to facilitate P_i_ scavenging from the environment, low-affinity transporters are degraded and replaced by high-affinity transporters, such as Pho84 (18–21). Under complete P_i_ starvation, when P_i_ can no longer be obtained from the environment, yeast adopts different strategies. Cells recycle P_i_ by decomposing various intracellular organic molecules including lipids, nucleotides, and even subcellular organelles (22–26). In addition, cells stop the cell cycle, saving P_i_ that would otherwise be used for nucleic acid and phospholipid duplication (27, 28).

In addition to scavenging and recycling, yeast cells also maintain a large store of phosphate inside their vacuoles (29, 30). These vacuoles share key features of acidocalcisomes, a class of conserved lysosome-like organelles that occur in all eukaryotic kingdoms (31, 32). These features include high lumenal concentrations of divalent cations, inorganic polyphosphates (polyPs), and basic amino acids. Their high polyP content is of potential relevance to phosphate homeostasis (14, 33, 34). This polymer can accumulate to the equivalent of hundreds of millimolar of phosphate units in an osmotically almost inert form and is considered as an efficient storage form of P_i_ (30, 35). Vacuolar polyP is produced from cytosolic ATP by the vacuolar transport chaperone (VTC) complex, which at the same time translocates the nascent polyP chain into the lumen of the organelle (36–38). PolyP can be hydrolyzed by polyphosphatases in the lumen of the organelle (39–42). It is assumed that the liberated P_i_ is released back into the cytosol to support metabolism. Therefore, polyP was proposed to act as a buffer system for cytosolic P_i_ (14, 43, 44). Synthesis of polyP by VTC is stimulated by inositol pyrophosphates (5, 45, 46). The levels of all these compounds (in yeast: 1-InsP_7_, 5-InsP_7_ and 1,5-InsP_8_) increase in abundance with increasing P_i_ availability (Chabert et al., manuscript in preparation)(47) and will hence favor accumulation and storage of P_i_ in the form of polyP only if P_i_ is abundant. How the turnover of polyP inside the organelle is regulated and coordinated with the release of P_i_ into the cytosol is unknown.

To explore mechanisms that contribute to an early recognition of forthcoming P_i_ limitation we explored the metabolic consequences experienced by yeast cells that are at the brink of phosphate limitation, i.e. where P_i_ becomes scarce but not yet limiting for growth. Furthermore, we analyzed the metabolic significance of acidocalcisome-like vacuoles and their polyP pool.

## Results

### Optimization of P_i_-starvation conditions for metabolomic analysis

Yeast cells distinguish scarcity of P_i_, where they induce genes to optimize P_i_ scavenging but can maintain normal growth, from P_i_ starvation that reduces growth and induces genes to facilitate P_i_ recycling from internal sources (6, 14–16, 20, 48, 49). We used non-targeted metabolomic analysis to explore whether and how cells react to beginning P_i_ scarcity in comparison to profound P_i_ starvation. To this end, we first identified conditions that bring our strain background to the brink of P_i_ limitation of growth. Batch cultures of cells were grown logarithmically overnight. A small inoculum of these logarithmically growing cells was transferred into synthetic complete (SC) liquid medium containing 0-10 mM P_i_ and the optical density (OD_600_) was measured for 24 h. Compared to cells growing in high-P_i_ medium (10 mM P_i_) growth was normal at 1 mM and 0.5 mM P_i_, and we observed only a mild retardation of growth in 0.25 mM and 0.1 mM P_i_. Incubation in 0 mM P_i_ arrested growth (Figure 1A). We measured the activity of secreted acid phosphatase, which is activated by P_i_ starvation (17), as a read-out of the PHO pathway. In P_i_-free medium, the activity of secreted acid phosphatase gradually increased up to 8 h and then maintained a similar level up to 24 h, suggesting that the PHO pathway was maximally activated at 8 h (Figure 1B). In 0.1 mM and 0.25 mM P_i_, the activity of acid phosphatase was partially induced at 8 h but fully induced at 24 h, probably reflecting the gradual depletion of P_i_ from the medium over that period. In 0.5 mM P_i_ media, acid phosphatase activity increased to half of the maximal value on 0 mM P_i_. There was no significant growth delay under this condition, indicating that the cells managed to compensate the limited availability of P_i_, probably through induction of the PHO pathway (Figure 1A). We monitored this induction by GFP-tagging the Pho4 transcription factor, which is translocated into the nucleus when cells lack P_i_. Pho4-GFP was predominantly in the nucleus after 8 h of incubation without P_i_ (Figure 1C). 0.5 mM P_i_ led to partial nuclear accumulation of Pho4-GFP, suggesting that the PHO pathway was moderately activated. We further measured the transcript level of the PHO pathway marker genes Pho5 and Pho84 using quantitative real-time reverse transcription PCR. The gene expression of Pho5 and Pho84 was highly induced after 8 h of P_i_ starvation (Figure 1D and E). Under P_i_ scarcity (0.5 mM P_i_), however, only Pho84 showed a mild yet statistically significant induction, whereas induction of Pho5 remained below 1 % of the maximal value and was not significant. In agreement with earlier studies (14, 50), this suggests that partial induction of the PHO pathway allows the cells to compensate for reduced availability of P_i_ to a level that supports normal growth.

**Figure 1.**
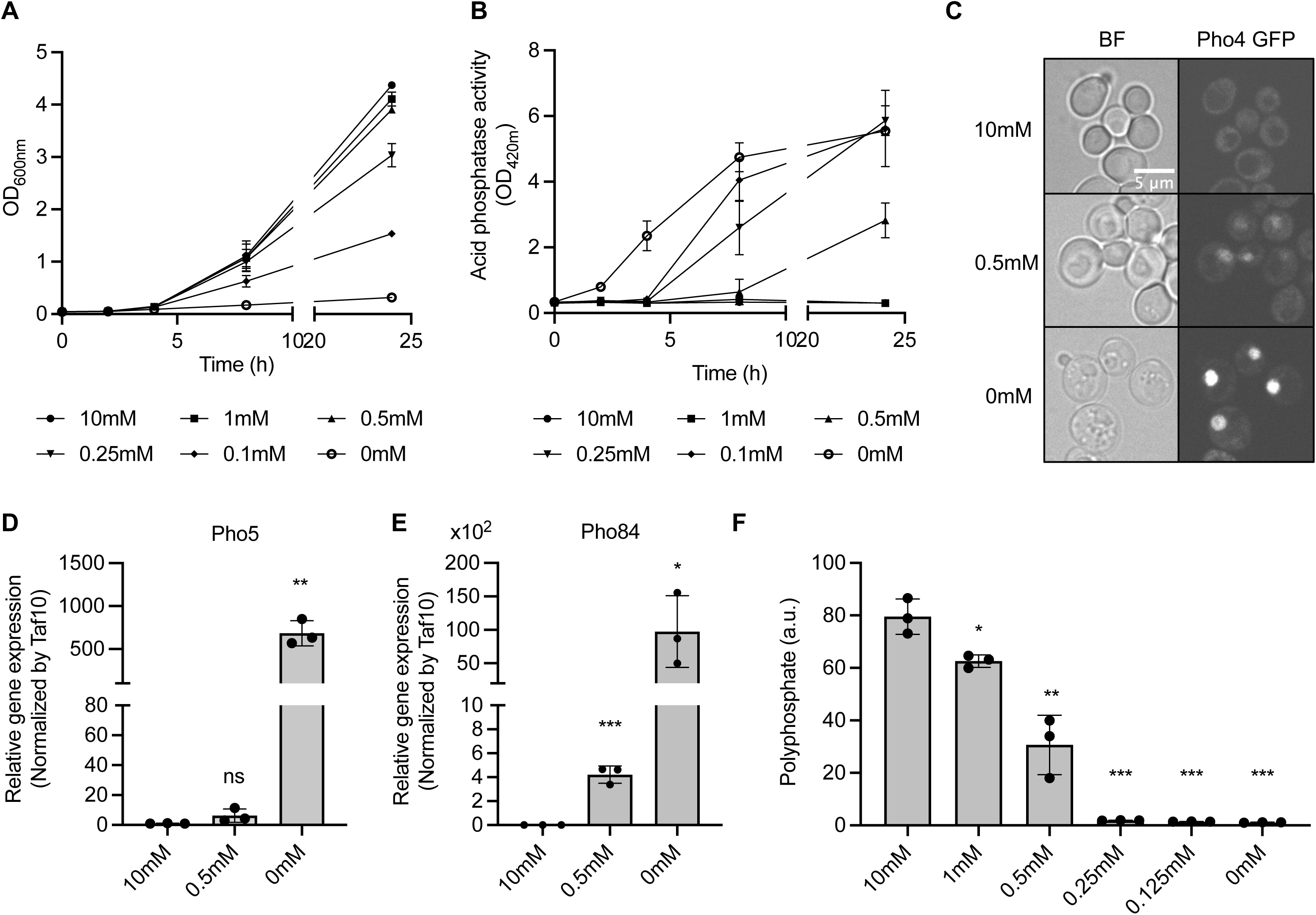
Response of *S. cerevisiae* under different Pi starving conditions. **A.** Growth curves of yeast cells in synthetic complete media supplemented with different concentrations of Pi from 10 mM to 0 mM. Cells were inoculated at an OD_600_ of 0.05 and cultured for 24 h. The means of triplicates are shown with standard deviation. **B.** Acid phosphatase activities of yeast cells grown as in A. The means of triplicates are shown with standard deviation. **C**. Fluorescence microscopy of live yeast cells expressing Pho4 genomically tagged with GFP. Cells were incubated for 8 h in 10 mM, 0.5 mM, and 0 mM Pi media as in A before observation. **D** and **E**. Relative gene expression levels of Pho5 (D) and Pho84 (E). Cells were grown in 10 mM, 0.5 mM, and 0 mM Pi media for 8 h and harvested for qRT-PCR. Fold changes were normalized with internal control Taf10. The means of three biological replicates are shown with standard deviation. ***, p < 0.001; **, p < 0.01; *, p < 0.05 by Student’s t-test. **F**. Polyphosphate levels in different Pi-containing media. Cells were incubated for 8 h as in A and harvested for polyphosphate measurement. The means of triplicates are shown with standard deviation. ***, p < 0.001; **, p < 0.01; *, p < 0.05; ns, not significant by Student’s t-test.

Under P_i_-limiting conditions, vacuolar polyP is degraded (14, 30, 41, 42, 51). It is assumed that resulting P_i_ is exported from the vacuoles to replenish the cytosolic P_i_ pool. In 0.25 mM, 0.125 mM and 0 mM P_i_, polyP was completely degraded (Figure 1F). However, after 8 h of growth in 0.5 mM P_i_, cells retained 40% of polyP compared to P_i_-replete conditions. We hence chose the described scheme of 8 h of growth in 0.5 mM P_i_ as a condition for metabolomic analysis under P_i_ limitation, because it partially activates the PHO pathway, has no significant effect on cell growth, and allows partial maintenance of the vacuolar polyP pool.

### Complete P_i_ starvation induces broad metabolic changes

For metabolome analyses logarithmically growing cells were transferred into media providing abundant P_i_ (10 mM), P_i_-scarcity (0.5 mM) or P_i_-starvation (0 mM P_i_). After 8 h of incubation in these media, 0.5 OD_600nm_ units of cells were harvested using vacuum filtration through a polytetrafluoroethylene polymer (PTFE) membrane and immediately frozen in liquid nitrogen (52). To analyze the metabolic effects of P_i_ limitation, untargeted metabolomic analyses were conducted through hydrophilic interaction liquid chromatography-mass spectrometry (HILIC-MS). A total of 169 metabolites were identified. Individual metabolic features were normalized by the median of each sample, transformed to log_10_, and centralized to the mean by an autoscaling method for further statistical analyses. Partial least squares discriminate analysis (PLS-DA) was performed to condense the metabolomic data into a simple plot allowing easy comparison of overall metabolic features (Figure 2A). This revealed a clear separation of the three growth conditions. Component 1, which comprises the largest difference of the total variance in metabolites (53.3%), placed the 0 mM P_i_ samples far from 10 mM and 0.5 mM P_i_ samples, suggesting that the metabolic changes caused by P_i_ starvation were greater than by P_i_ limitation (Figure 2A). In the loading plot, which visualizes the contribution of individual metabolites to components 1 and 2, most phosphate-containing metabolites (red opened circle) negatively contributed to component 1, indicating that they decreased under P_i_-starvation (Figure 2B). Only two phosphate-containing metabolites, AICAR and 3’,5’ cyclic GMP, positively contributed to component 1. For further analysis, we compared variable importance in projection (VIP) scores of each metabolite, which represent the contribution of variables to the PLS-DA model encompassing 10 mM, 0.5 mM and 0 mM P_i_ data. ATP showed the highest VIP value among the top 20 most influential metabolites (Figure 2C), underscoring an interrelation of P_i_ and ATP metabolism (Figure 2C)(53).

**Figure 2.**
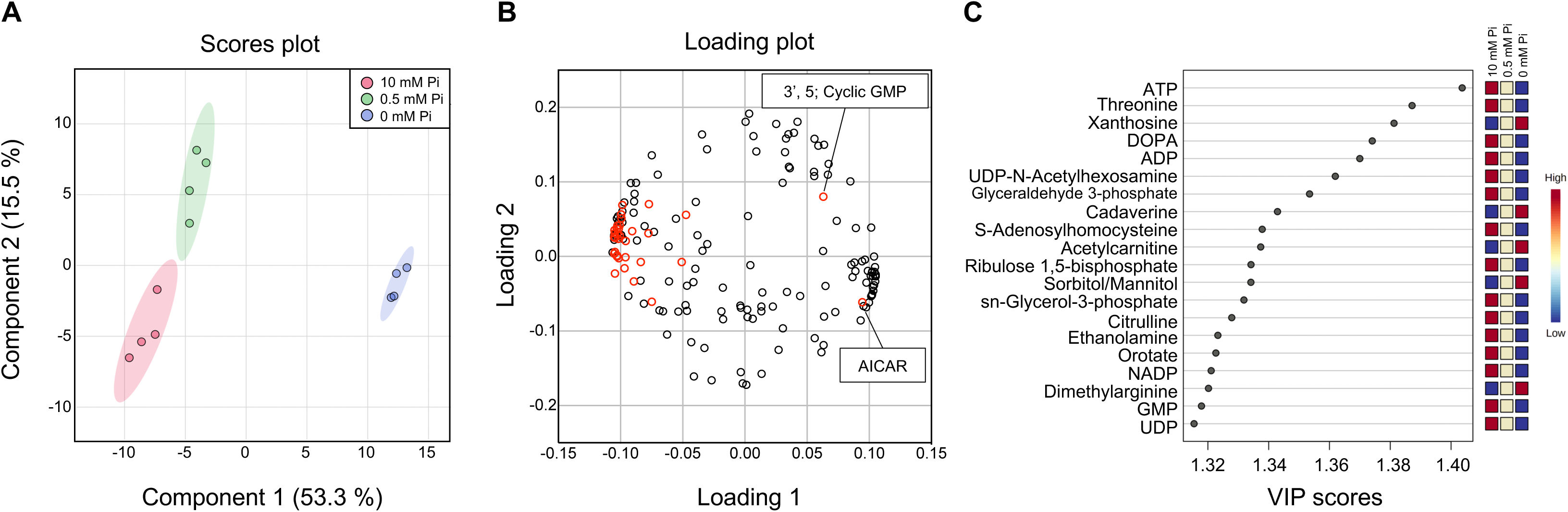
Partial least squares discriminant analysis (PLS-DA) of *S. cerevisiae* metabolites under different Pi conditions. **A.** Score plot of PLS-DA. Red, green, and blue dots indicate the replicates of yeast metabolomic data incubated in 10 mM, 0.5 mM, and 0 mM Pi media, respectively. The shaded regions represent the 95% confidence intervals. **B**. Loading plot of PLS-DA. Red dots indicate Pi-containing metabolites. **C**. Variable of importance in projection (VIP) scores of the top 20 metabolites generated from PLS-DA. The color code indicates the relative abundance of each metabolites in different Pi conditions.

Pearson’s coefficients were calculated as a measure for the correlation between metabolites. Two metabolic groups (group 1 and group 2) were negatively correlated with each other (Figure 3A). The relative abundance of metabolites in 0 mM P_i_ medium increased for group 1 and decreased for group 2 (Figure 3B and C). Purine and pyrimidine pathway metabolites, nucleosides and nucleobases increased (Table 1), mirrored by a decrease of nucleotides. Metabolites of the citrate cycle increased, such as citric acid, isocitric acid, and oxoglutaric acid, whereas the levels of glycolytic intermediates declined, suggesting an altered strategy for energy production under P_i_-starvation. Nicotinate and nicotinamide metabolites also decreased. By contrast, metabolites of the tryptophan degradation pathway, which are involved in NAD *de novo* synthesis, accumulated.

**Figure 3.**
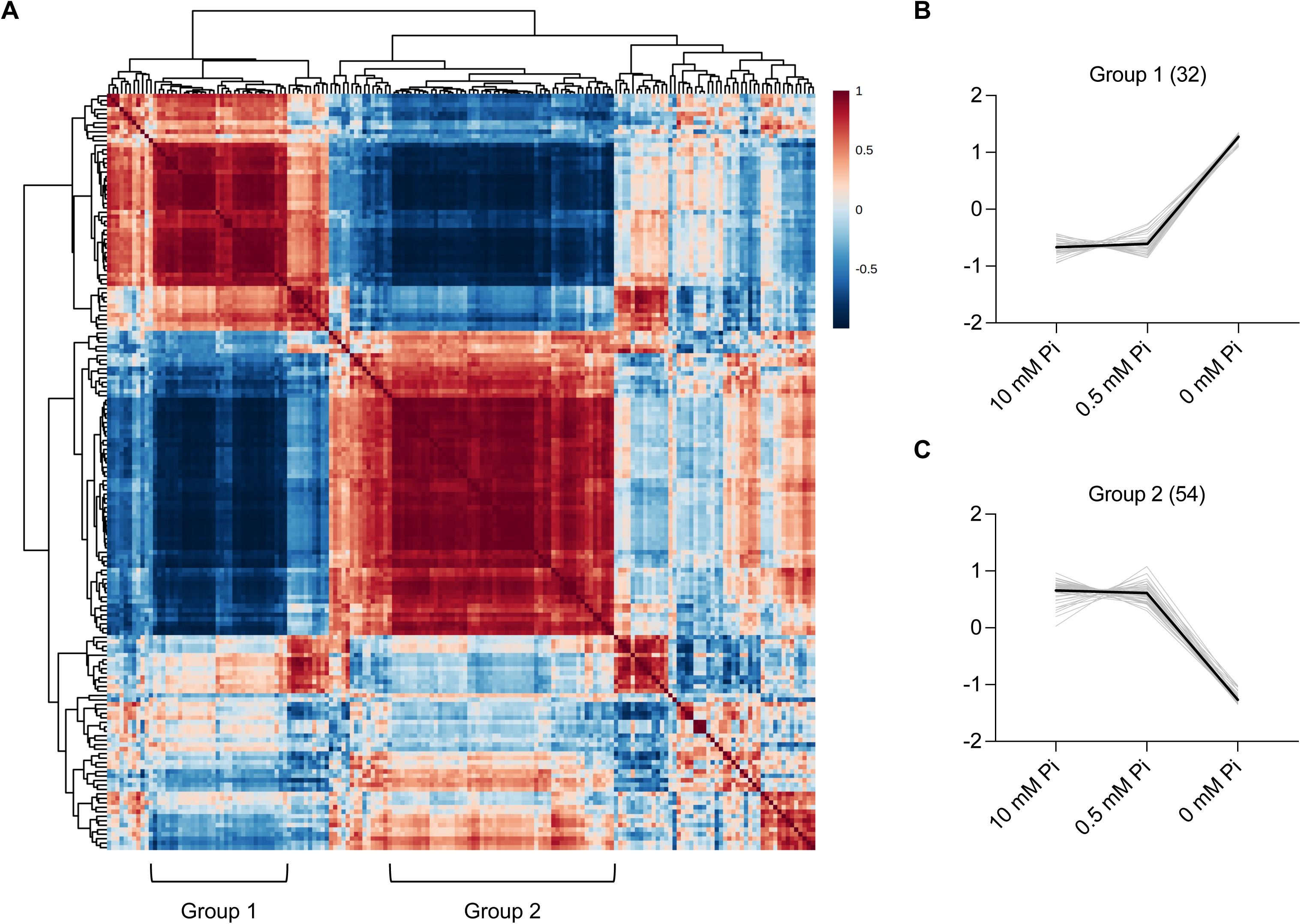
Correlation analysis of *S. cerevisiae* metabolites under different Pi starving conditions. **A.** Clustered correlation heat-map between metabolites under different Pi conditions. The correlation matrix was generated by Pearson correlation coefficients, which are represented by a color code. Red and blue indicate positive and negative correlations, respectively. Two representative metabolic groups showing strong correlations are marked as group 1 and group 2. **B** and **C**. Relative abundance of metabolites included in group 1 (B) and group 2 (C) under different Pi conditions. The profiles of individual metabolites are shown in grey.

**Table 1.** List of metabolites in group 1 and group 2 from the correlation heatmap.

|  | Metabolic pathway | Metabolites |
| --- | --- | --- |
| Group 1 | Purine | Deoxyadenosine, Adenosine<br>AICAR, Guanosine, Inosine, Xanthine, Xanthosine |
|  | Pyrimidine | Cytosine, Uridine |
|  | TCA | Citric acid, Isocitric acid Oxoglutaric acid |
|  | Cysteine and methionine | S-Adenosylmethionine, 5'-Methylthioadenosine |
|  | Tryptophan | Kynurenine, Kynurenic acid, Xanthurenic acid, Quinolinic acid |
|  | Glycerolipid | Choline, Glyceric acid |
|  | Others | 5'-Deoxyadenosine, Acetylcarnitine, Cadaverine, Glutamine, Methylglutaric acid, N-Acetylphenylalanine, N-Methylglutamic acid, N-6-Trimethyllysine, p-Hydroxyphenylacetic acid, Pantothenic acid, Pyridoxal |
| Group 2 | Purine | AMP, ADP, ADP ribose, ATP, dATP, GMP, GDP, Guanine, IMP, Hypoxanthine |
|  | Pyrimidine | CMP, CDP, dCDP, UMP, UDP, Orotic acid |
|  | Glycolysis | Acetyl-CoA, Fructose 6-phosphate, Glyceraldehyde 3-phosphate, Glucose 6-phosphate, 2/3-Phosphoglyceric acid, Phosphoenolpyruvic acid |
|  | Cysteine and methionine | 1-Aminocyclopropanecarboxylic acid, Ophthalmic acid, S-Adenosylhomocysteine, Cystathionine, 3-Sulfinoalanine, Pyroglutamic acid |
|  | Nicotinate and nicotinamide | NAD<sup>+</sup>, NADP<sup>+</sup>, Nicotinic acid mononucleotide |
|  | Glycerolipid | sn-Glycerol-3-phosphate, Ethanolamine, Glycerol |
|  | Arginine | N-Acetylputrescine, Proline, N-Acetylglutamic acid, Citrulline, Dimethylarginine |
|  | Others | Guaiacol, Aminoisobutyric acid, 2-Hydroxyglutaric acid, Aspartic acid, Glutamic acid, Glycine, Histidine, Lysine, Serine, Threonine, Mevalonic acid, N(2)-Acetyllysine, TMP, UDP-hexose, UDP-N-Acetylhexosamine |

### Pathway analysis of P_i_ starvation

To identify metabolites that accumulated differentially and in a statistically significant manner, we compared their relative abundance by volcano plots. Profound metabolic effects occurred under complete P_i_ starvation: 31 metabolites increased and 49 decreased in a statistically meaningful way, representing 47% of the detected metabolites (Figure 4B, 4D). Pathway analysis of these 80 metabolites identified the most affected pathways as purine metabolism, pyrimidine metabolism, nicotinate and nicotinamide metabolism, glycolysis/gluconeogenesis, citrate cycle, and cysteine and methionine metabolism, with - log(p)>1.5 (Figure 5B and Table 2).

**Figure 4.**
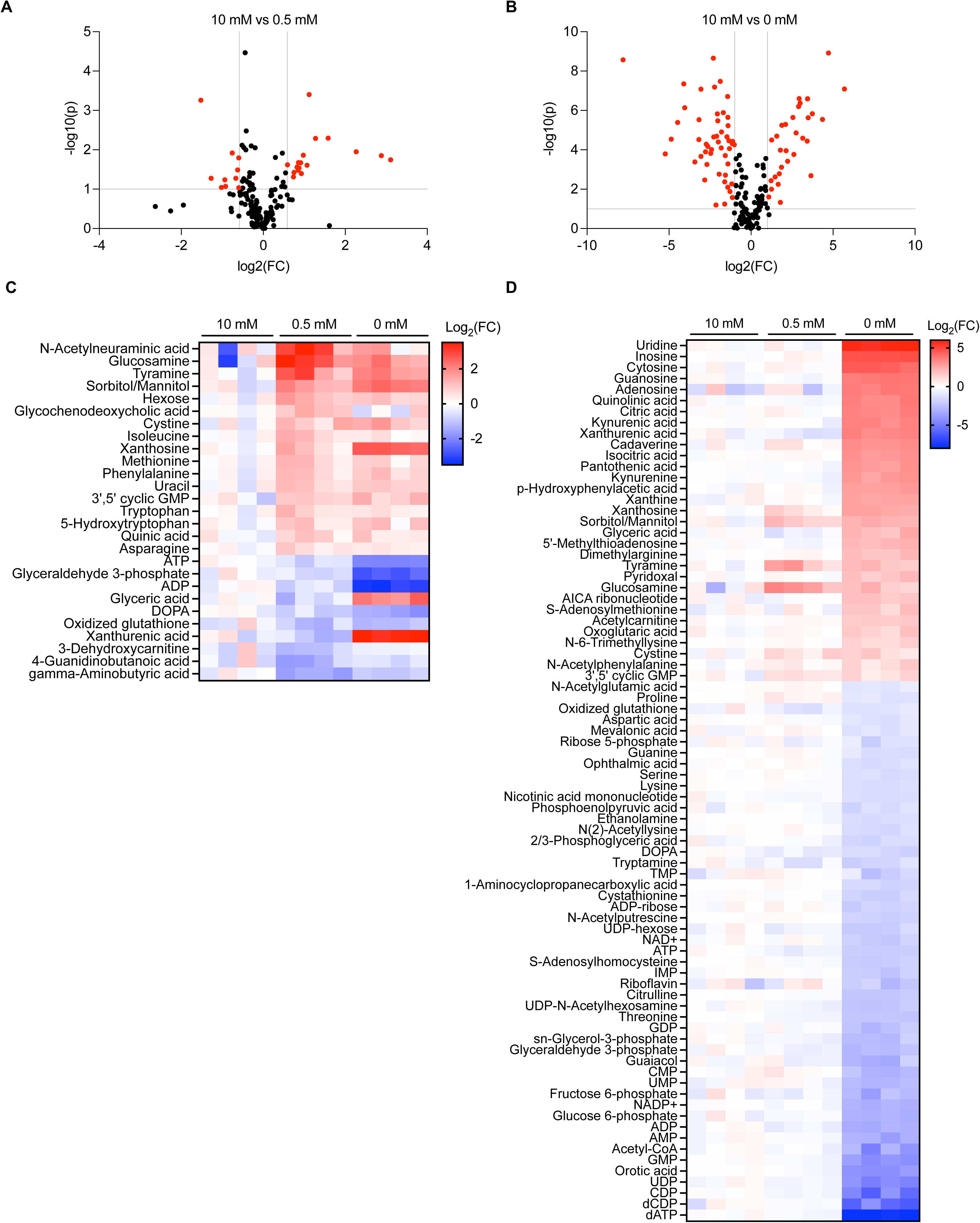
Metabolites differentially accumulated under Pi scarcity or Pi starvation. **A** and **B**. Volcano plot analysis of changes upon Pi scarcity and Pi starvation. (A) Changes upon Pi scarcity (0.5 mM Pi). Red dots indicate differentially accumulated metabolites (fold change, |FC| > 1.5; p < 0.1. (B) Changes upon Pi starvation (0 mM Pi). Red dots indicate differentially accumulated metabolites (fold change, |FC| > 1.5; p < 0.1). These metabolites are listed in Supplementary Tables 1 and 2. (C) Heat map of Pi scarcity. The list was selected by t-test (p < 0.1), showing metabolites changing at least 1.5-fold under Pi scarcity (0.5 mM Pi). (D) Heat map of Pi starvation. Same as in C but showing metabolites changing at least 2-fold under Pi starvation (0mM Pi). The relative abundance of metabolites is represented as log_2_(fold-change) through a color code.

**Figure 5.**
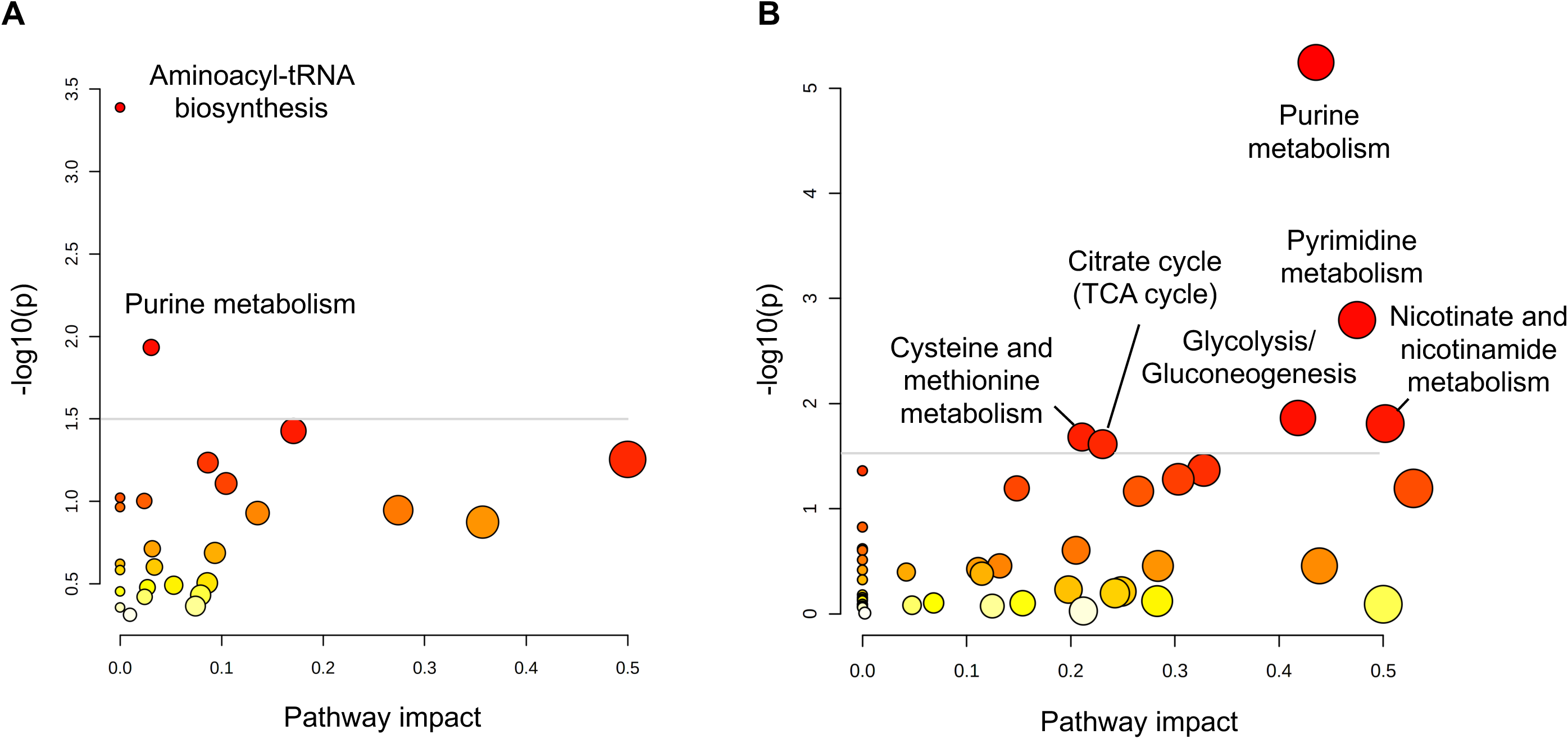
Pathway analysis of differentially accumulated metabolites under different Pi regimes. **A** and **B**. Pathway analysis of differentially accumulated metabolites upon Pi scarcity (0.5 mM Pi) (A) and Pi starvation (0 mM Pi) (B). The size and color of the circle indicate impact value and p-value, respectively. The annotated metabolic pathways have higher statistical significance (-log(p) > 1.5).

**Table 2.** The list of top 15 metabolic pathways differentially accumulated under P_i_ starvation.

| Pathway | Match status | p-value | -log(p) | Impact |
| --- | --- | --- | --- | --- |
| Purine metabolism | 17/62 | 0.0000056885 | 5.245 | 0.43543 |
| Pyrimidine metabolism | 9/34 | 0.0016028 | 2.7951 | 0.47488 |
| Glycolysis or gluconeogenesis | 6/24 | 0.013697 | 1.8634 | 0.41811 |
| Nicotinate and nicotinamide metabolism | 4/12 | 0.015489 | 1.81 | 0.50185 |
| Cysteine and methionine metabolism | 8/41 | 0.02079 | 1.6822 | 0.21078 |
| Citrate cycle (TCA cycle) | 5/20 | 0.024305 | 1.6143 | 0.2306 |
| Methane metabolism | 5/23 | 0.042866 | 1.3679 | 0.32774 |
| Lysine biosynthesis | 4/16 | 0.043511 | 1.3614 | 0.0 |
| Glycine, serine and threonine metabolism | 6/32 | 0.052435 | 1.2804 | 0.30324 |
| Arginine biosynthesis | 4/18 | 0.064018 | 1.1937 | 0.14819 |
| Pentose phosphate pathway | 4/18 | 0.064018 | 1.1937 | 0.52896 |
| Glyoxylate and dicarboxylate metabolism | 5/26 | 0.068182 | 1.1663 | 0.26506 |
| Cyanoamino acid metabolism | 2/8 | 0.14937 | 0.82572 | 0.0 |
| Synthesis and degradation of ketone bodies | 1/3 | 0.23995 | 0.61987 | 0.0 |
| Beta-alanine metabolism | 2/11 | 0.24839 | 0.60487 | 0.0 |

#### Glycolysis & TCA cycle

P_i_ starvation reduced the abundance of glycolysis intermediates 2- to 10-fold, including glucose-6-phosphate, fructose-6-phosphate, glyceraldehyde 3-phosphate, 2/3 phosphoglyceric acid, phosphoenolpyruvic acid and acetyl-CoA (Supplementary Figure 1). In addition, ribose 5-phosphate, which is derived from glucose 6-phosphate via the oxidative pentose phosphate pathway, was reduced. The upstream TCA cycle metabolites including citric acid, isocitric acid, and oxoglutaric acid accumulated 2- to 10-fold, whereas later TCA cycle metabolites, including succinic acid and malic acid, did not change (Supplementary Figure 2). Thus, P_i_ starvation has a strong impact on glycolysis and the TCA cycle.

#### Nicotinate and nicotinamide metabolism

The P_i_-containing metabolites nicotinic acid mononucleotide, NAD^+^, and NADP^+^ decreased 2- to 5-fold under P_i_ starvation (Supplementary Figure 3). In contrast, metabolites of NAD^+^ *de novo* synthesis, also known as the kynurenine pathway, accumulated more than 6-fold. Based on these results, it can be speculated that the impaired NAD^+^ synthesis from the kynurenine pathway will affect intracellular redox homeostasis..

#### Cysteine and methionine metabolism

S-adenosylmethionine (SAM) is the major methyl donor for modifications of various biomolecules including proteins, DNA, RNA, and metabolites, producing S-adenosylhomocysteine (SAH) as a byproduct of the reaction. The relative abundance of SAM and of methionine salvage pathway metabolites increased, while that of SAH decreased under P_i_-starvation (Supplementary Figure 4). Cystathionine, which can be produced from SAH via homocysteine, diminished under P_i_ starvation, whereas metabolites derived from cystathionine, such as glutathione and taurine, were not significantly affected. P_i_ starvation also affected another SAM-dependent branch, the synthesis of polyamines, because a byproduct of polyamine synthesis, 5’-methylthioadenosine (MTA), accumulated strongly. These results suggest that yeast cells change their strategy of SAM utilization under P_i_-deprivation. The reduction of nucleic acid synthesis, which accompanies growth arrest, may reduce consumption of SAM for nucleotide synthesis and promote accumulation of this compound.

#### Purine and pyrimidine metabolism

Nucleosides and nucleobases such as adenosine, inosine, xanthosine, guanosine, uridine, cytosine, and xanthine significantly accumulated under P_i_ starvation (Supplementary Figure 5 and 6) whereas P_i_-containing nucleotides, nucleoside diphosphates, and nucleoside triphosphates, decreased. By contrast, the amount of AICAR increased. This metabolic change may contribute to triggering the PHO pathway under P_i_ starvation in two ways. First, AICAR inhibits the production of InsP_8_ (54), which itself is a potent suppressor of the PHO pathway (Chabert et al., in preparation). Second, AICAR stabilizes the interaction of the association of the transcription factors Pho4 and Pho2, which is necessary for full induction of the PHO pathway (13).

### Moderate P_i_ limitation causes few, potentially diagnostic metabolic changes

Few significant changes (-log(p)>1) occurred under P_i_ limiting conditions: Only 17 metabolites increased and 10 decreased more than 1.5-fold (Figure 4A, 4C, 5A). ATP decreased in a statistically significant manner, which suggests that the ATP level is more sensitive to P_i_ availability than that of most other metabolites, making ATP a bona fide early indicator of P_i_ scarcity. In line with this, key enzymes for signaling the intracellular P_i_ state, InsP_6_ kinases, have an unusually high Km for ATP, which is close to the normal cellular concentration of this compound (55, 56). To test this possibility, we measured the products of these enzymes, inositol pyrophosphates, under different P_i_ conditions using capillary electrophoresis coupled electrospray ionization mass spectrometry (CE-ESI-MS). Under P_i_ starvation, 1,5-InsP_8_ was not detected at all and 5-InsP_7_ and 1-InsP_7_ decreased by 80% (Figure 6). Even under mild P_i_ limitation, all three compounds significantly declined in comparison with P_i_-replete conditions, by 60% for 1,5-InsP_8_, 30% for 5-InsP_7_, and 50% for 1-InsP_7_. This is consistent with the hypothesis that even moderate decreases in ATP levels under P_i_ limitation can be translated into decreased InsPP levels, which then activate SPX-domain-based signaling to stabilize cytosolic P_i_.

**Figure 6.**
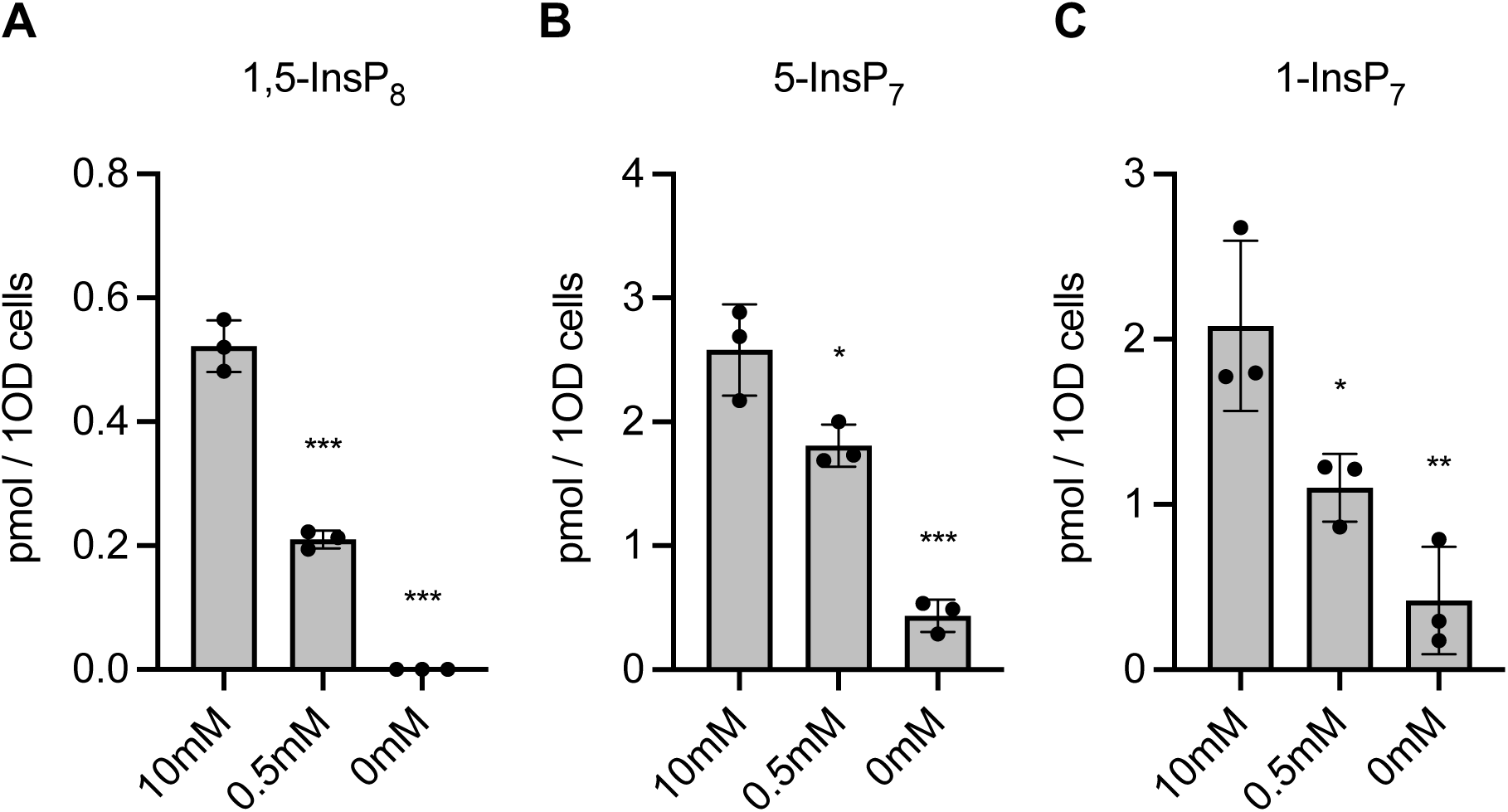
Inositol pyrophosphates profiles under different Pi conditions. **A-C.** Inositol pyrophosphates levels of yeast cells grown in 10 mM, 0.5 mM, and 0 mM Pi media for 8 h. The data was normalized by OD_600_. The means of triplicates are shown with standard deviation. ***, p < 0.001; **, p < 0.01; *, p < 0.05 by Student’s t-test.

### PolyP in acidocalcisome-like vacuoles contributes to P_i_ homeostasis even in P_i_-replete conditions

Yeast cells contain acidocalcisome-like vacuoles, which can convert the γ-phosphate from ATP into inorganic polyphosphate. Thereby they can store hundreds of millimolar of phosphate units in an osmotically inactive form (36, 37). At the same time, vacuoles contain polyphosphatases and P_i_ exporters (42). This system is powerful enough to influence P_i_ homeostasis of the cells and the P_i_ starvation response when it is dysregulated (14, 33, 34, 51). To investigate the metabolic role of vacuolar polyphosphates, we analyzed the global metabolic profiles of the vtc4Δ mutant, which lacks the polyP-synthesizing complex VTC, under P_i_-replete conditions and after 2 hours of P_i_ starvation. A PLS-DA plot showed that the metabolic features of vtc4Δ cells were clearly distinct from that of the wildtype, in P_i_-rich conditions and even more under P_i_ starvation (Figure 7A). The loading plot of PLS-DA revealed that almost all phosphate-containing metabolites (red opened circle) were abundant in P_i_-rich conditions, except AICAR, consistent with the results above (Figure 7B and Figure 2B).

**Figure 7.**
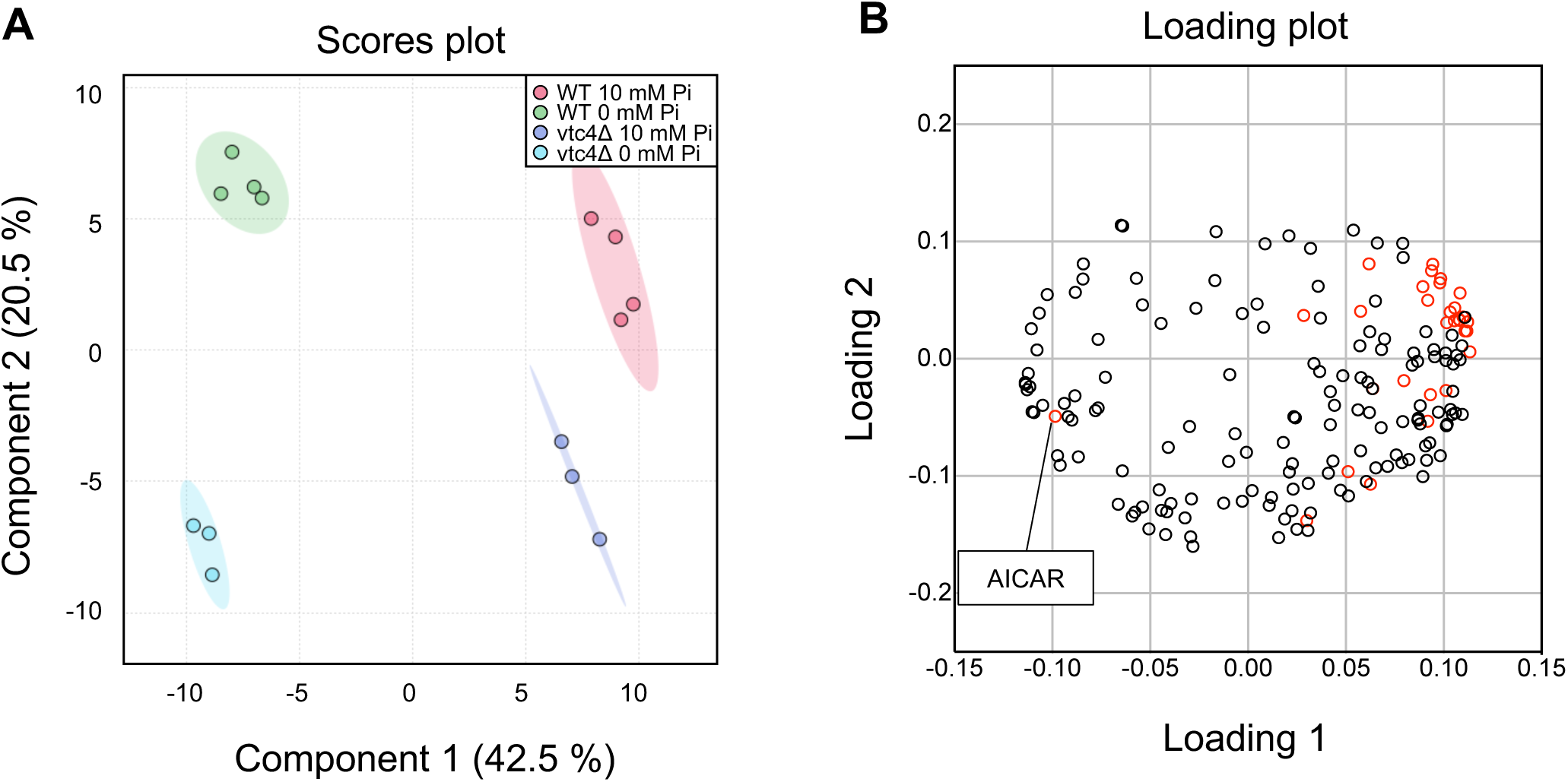
Multivariate statistical analysis of metabolite profiling data from wild type and vtc4Δ under 10 mM and 0 mM Pi conditions. **A.** Score plot of partial least squares-discriminant analysis (PLS-DA). Red, green, blue, light blue indicate the replicates of metabolomic data from wild type 10mM Pi, wild type 0 mM Pi, vtc4Δ 10 mM Pi, and vtc4Δ 0 mM Pi, respectively. The shaded regions represent the 95% confidence intervals. **B**. Loading plot of PLS-DA. Red dots indicate Pi-containing metabolites.

To analyze how polyphosphate synthesis and P_i_ concentration in the medium affect metabolic features, a two-way ANOVA test was performed. The relative abundance of 92, 116, and 78 metabolic features were affected by polyP, P_i_ concentration, and their interaction, respectively (Figure 8A). A total of 65 metabolites were affected by both polyP and P_i_, of which 66 % (43 features) were additionally affected by their interaction. Metabolite set enrichment analysis was performed using these 43 metabolites. Most metabolic pathways affected by P_i_ starvation shown in the previous pathway analysis, such as pyrimidine metabolism, nicotinate and nicotinamide metabolism, glycolysis, and purine metabolites, were again ranked statistically high, showing consistency of the analyses (Figure 8B and Table 3). The changes of these 43 metabolites were visualized by heat-map analysis (Figure 8C). vtc4Δ cells showed much more pronounced changes than wildtype cells in several respects: The decrease of P_i_-containing purines and pyrimidines (CMP, UMP, AMP, and dGMP) (Figure 8C, Supplementary Figure 7 and 8); the increase of nucleosides and nucleobases (cytosine, cytidine, guanosine, uridine, and inosine) (Figure 8C, Supplementary Figure 7 and 8); and the reduction of NAD^+^ and NADP^+^ (Supplementary Figure 9). vtc4Δ cells grown on P_i_-replete medium showed numerous metabolic features of P_i_-starved wildtype cells. They did not, however, simply phenocopy a P_i_ starvation response at a reduced scale. Intermediates of tryptophan degradation, such as kynurenine, kynurenic acid, 3-hydroxykynurenine, and the NAD+ precursor quinolinic acid, which did not change significantly in P_i_ starved wildtype cells, were increased in vtc4Δ already under P_i_-replete conditions and increased further (up to 5-fold) upon P_i_ starvation (Figure 8C and Supplementary Figure 9). In vtc4Δ cells, the later glycolytic intermediates dihydroxyacetone phosphate and phosphoenolpyruvate were more abundant than in the wildtype in P_i_ replete medium, and 2/3 phosphoglyceric acid underwent a much more pronounced reduction than in the wildtype upon P_i_ starvation (Figure 8C and Supplementary Figure 10). These results indicate that polyP synthesis by VTC has a significant effect on the metabolic profile of cells. It dampens the metabolic consequences of P_i_ starvation, which is consistent with its proposed role as a P_i_ reserve. But it also has so far unrecognized metabolic functions under P_i_-replete conditions, as indicated by metabolic features of vtc4Δ cells that cannot be recapitulated by P_i_ scarcity or P_i_ starvation of wildtype cells.

**Figure 8.**
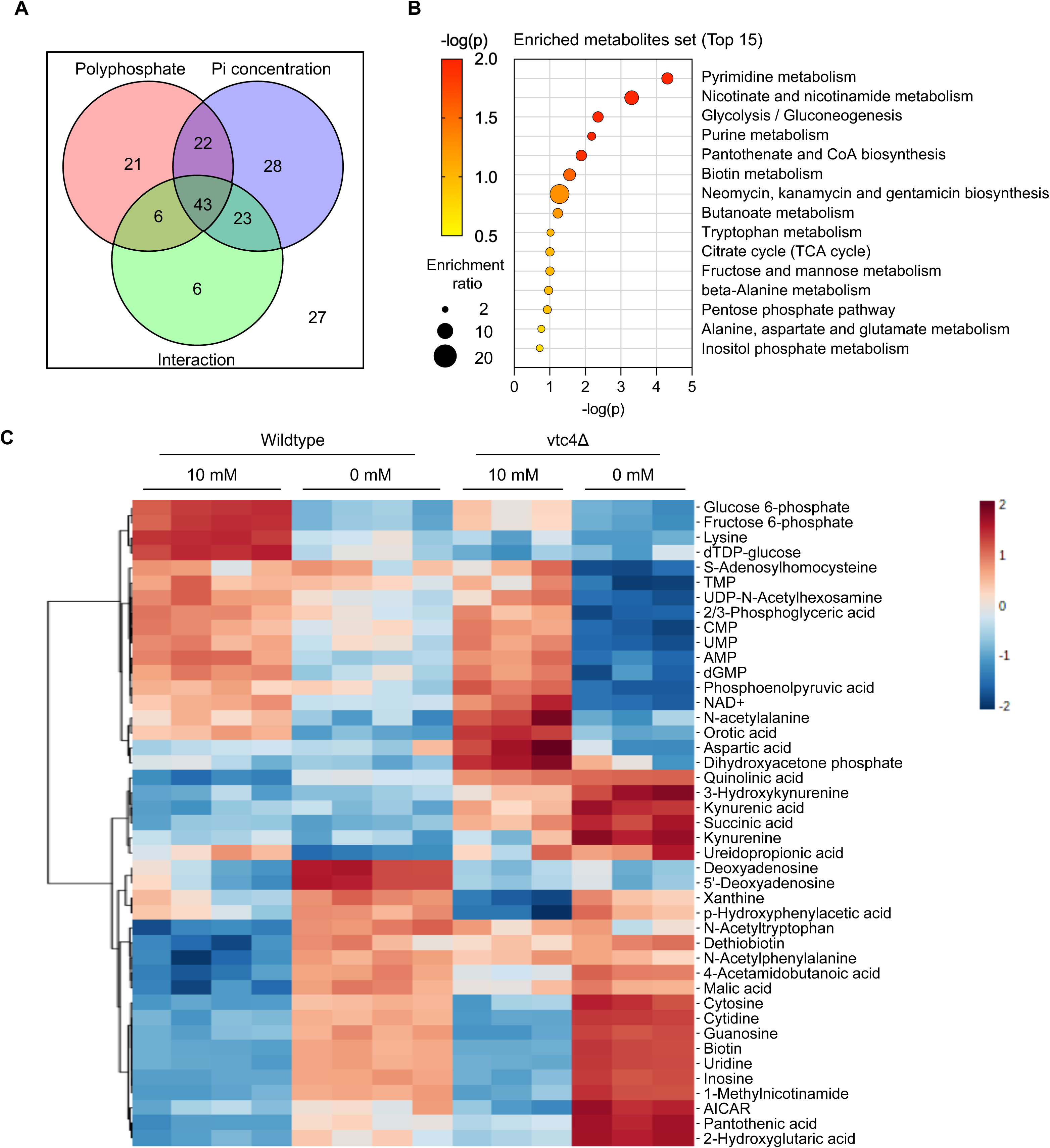
Interrelated effect of polyphosphate and Pi starvation on metabolic pathways. **A.** Summary of two-way ANOVA analysis (adj p-value < 0.05). Red, blue, and green represent the metabolites affected by polyphosphate (wild-type and vtc4Δ), Pi concentration (10 mM and 0 mM Pi), and interaction between both (polyphosphate and Pi concentration), respectively. **B**. Metabolite set enrichment analysis of 43 metabolites simultaneously affected by polyphosphate, Pi concentration, and their interaction. The top 20 metabolite sets were selected based on p-value. **C**. Heatmap was generated based on the list of 43 metabolites affected by Pi concentration, polyP and their interaction from a two-way ANOVA. Features were clustered by Euclidean distance using Ward’s clustering method. The color code indicates the normalized intensity of metabolic features

**Table 3.** The top 15 metabolite set from metabolite set enrichment analysis.

| Metabolite set | Total | Hits | Expected | p-value |
| --- | --- | --- | --- | --- |
| Pyrimidine metabolism | 39 | 7 | 1.05 | 0.0000494 |
| Nicotinate and nicotinamide metabolism | 15 | 4 | 0.404 | 0.0005 |
| Glycolysis / Gluconeogenesis | 26 | 4 | 0.7 | 0.00441 |
| Purine metabolism | 65 | 6 | 1.75 | 0.00666 |
| Pantothenate and CoA biosynthesis | 19 | 3 | 0.512 | 0.013 |
| Biotin metabolism | 10 | 2 | 0.269 | 0.0278 |
| Neomycin, kanamycin and gentamicin biosynthesis | 2 | 1 | 0.0539 | 0.0532 |
| Butanoate metabolism | 15 | 2 | 0.404 | 0.0596 |
| Tryptophan metabolism | 41 | 3 | 1.1 | 0.0954 |
| Citrate cycle (TCA cycle) | 20 | 2 | 0.539 | 0.0991 |
| Fructose and mannose metabolism | 20 | 2 | 0.539 | 0.0991 |
| beta-Alanine metabolism | 21 | 2 | 0.566 | 0.108 |
| Pentose phosphate pathway | 22 | 2 | 0.593 | 0.117 |
| Alanine, aspartate and glutamate metabolism | 28 | 2 | 0.754 | 0.173 |
| Inositol phosphate metabolism | 30 | 2 | 0.808 | 0.192 |

## Discussion

Our results extend earlier studies of P_i_ starvation (53, 57, 58). In agreement with these studies, we observed reductions in nucleotides and late glycolytic intermediates and increased nucleoside and nucleobase levels. These changes can be explained by simple mass action (58, 59) because they reduce phosphate-containing metabolites under conditions of intracellular P_i_ shortage. By contrast, early TCA cycle metabolites, such as citric acid, isocitric acid, and oxoglutaric acid, strongly increased during P_i_ starvation. Furthermore, oxygen consumption of yeast increases upon P_i_ starvation (60). This suggests that mitochondrial respiration may become activated as an alternative mechanism for energy production because it fixes less P_i_ in metabolic intermediates than ATP production based on glycolysis. In line with this, we made the side-observation that P_i_ starvation also caused mitochondrial fragmentation (data not shown). Mitochondria fragment in media favoring respiration, such as non-fermentable carbon sources or glucose-limited media (61, 62).

Upon P_i_ starvation, two P_i_-containing metabolites increased, AICAR and cyclic GMP (Figure 2B and Supplementary Figure 5). Little is known about the roles of cGMP in yeast so far (63). However, it provides a potential link to protein kinase A signaling, which is involved in P_i_ homeostasis and the signaling through P_i_ transporters in yeast (64–70). cGMP can also inhibit DNA polymerase (71), which might reduce P_i_ consumption by the cells and avoid P_i_ depletion during S-phase (43). AICAR is an intermediate of *de novo* purine biosynthesis. It activates a master regulator of energy homeostasis, AMP-activated protein kinase (AMPK) in mammalian cells (72) but the yeast AMPK Snf1 does not depend on it. Snf1 is activated by ADP (13, 73). Upon a block in nucleoside synthesis, accumulating AICAR stimulates the PHO transcription pathway by stabilizing the interaction of the responsible transcription factors Pho4 and Pho2 (13). However, it has remained unknown how AICAR behaves under P_i_ starvation. Our analysis now shows that AICAR accumulates during P_i_ starvation. AICAR accumulation may be favored by the decrease of ADP and ATP levels upon P_i_ starvation because these nucleotides exert feedback inhibition on the first step in purine synthesis and thereby on AICAR synthesis (74). Thus, AICAR may promote expression of PHO genes upon P_i_ starvation. However, under P_i_ scarcity, when ATP decreases less severely than under P_i_ starvation, AICAR did not increase. This corresponds to the only partial activation of the PHO pathway under P_i_ scarcity. We hence propose that an increase in AICAR may contribute to switching the PHO pathway from partial to full activation when cells transit from P_i_ scarcity to starvation.

Glucose 6-phosphate is used by the pentose phosphate pathway (PPP) to convert NADP^+^ to NADPH, which is essential for cellular redox homeostasis (75, 76). Although NADPH was not detected in our metabolomic analysis, NADP^+^ decreased, and we hence assume that NADPH should decrease as well. NAD^+^, nicotinic acid mononucleotide, and nicotinic acid, which are precursors of NADP^+^, were all reduced by P_i_ starvation, but intermediates of the kynurenine pathway, also known as the *de novo* NAD^+^ synthetic pathway, were significantly accumulated (Supplementary Figure 3). These changes may result from the accumulation of AICAR, which stimulates the expression of enzymes involved in the kynurenine pathway (77). The last metabolite of the kynurenine pathway, quinolinic acid, is converted to nicotinic acid mononucleotide by consuming phosphoribosyl pyrophosphate (PRPP). Since PRPP is produced from ribose 5-phosphate, this reaction may be impaired through the decrease in ribose-5-phosphate under P_i_ starvation, favoring the observed accumulation of quinolinic acid. In addition, the PHO pathway directly promotes the catabolism of NAD^+^ through inducing the vacuolar phosphatase Pho8, which removes P_i_ from nicotinic acid mononucleotide and nicotinamide mononucleotide (78). The decrease in the NAD^+^ and NADP^+^ pools is expected to affect intracellular redox homeostasis. This may increase the dependence of cells on the oxidative stress response for surviving Pi starvation, which had been observed (60). In line with this, cells with perturbed P_i_ and inositol pyrophosphate homeostasis induce the environmental stress response (79, 80).

We observed increased SAM and decreased SAH under P_i_ starvation. SAM is synthesized from methionine and ATP, releasing P_i_ and PP_i_. SAM provides methyl groups for methyl transfer reactions, generating SAH as a byproduct (81). Histone methylation affects global gene expression patterns by changing the structure of chromatin through interactions with various chromatin remodeling factors and transcription regulators (82, 83). The expression of PHO genes is also under the control of histone methylation: Expression of Pho5 and Pho84 is induced in the set1Δ mutant, which affects methylation of lysine 4 of the nhistone H3 (84, 85). In addition, the methyl transferase Hmt1 promotes expression of several P_i_-responsive genes (86). Thus, the changes of SAM upon P_i_ starvation might alter the intracellular methylation status and thereby provide a further route of input for P_i_-dependent gene expression.

An interesting question is how cells distinguish P_i_ scarcity from P_i_ starvation. This is challenging because P_i_ scarcity can be corrected by cells through partial induction of the PHO pathway. The resulting improved capacity for P_i_ scavenging, for example through expression of high-affinity P_i_ transporters, apparently allows them to maintain sufficient metabolic performance to support normal growth. Furthermore, positive feedback loops involving SPL2, a small regulator of the P_i_ transporter Pho90, may stably commit cells to the activation of the starvation program, even if this re-establishes sufficient intracellular P_i_ supply (16, 48, 49). Nevertheless, the P_i_ starvation program is not launched in a simple all-or-none fashion. Certain genes are activated at different levels of P_i_ shortage, as exemplified by the gene for the high-affinity transporter Pho84, which is activated earlier than that of the secreted acid phosphatase Pho5 (14–16). Our analyses of the low-P_i_ state, which were performed in cells that induced Pho84 but not Pho5, provide hints on metabolic changes that might be used by cells to distinguish P_i_ scarcity from P_i_ starvation. Under P_i_ scarcity, few metabolites changed in a statistically meaningful way, but the level of ATP and ADP decreased by 35% compared to P_i_-replete conditions (Supplementary Figure 5 and Supplementary Figure 7). Enzymes involved in the inositol pyrophosphate pathway have high K_m_ values for ATP (55, 56), so even a subtle decrease in ATP could significantly impair InsPP synthesis. In our experiments, the subtle decrease in ATP under P_i_ scarcity corresponded to a significant decrease in the InsPP pool (Figure 6). Since we recently observed that the PHO pathway is repressed by InsPPs through the SPX domain of Pho81 (Chabert et al., manuscript in preparation), we propose that this decline of InsPPs upon P_i_ scarcity inhibits Pho85/Pho80 activity, triggers partial Pho4 nuclear translocation and partial activation of the PHO pathway.

Yeast stores P_i_ in the form of polymers in acidocalcisome-like vacuoles. Under P_i_ limiting conditions, polyphosphatases degrade polyphosphate and liberate P_i_, which could potentially be brought back to the cytosol through vacuolar P_i_ transporter Pho91 (30, 41, 42). In this way, the vacuolar polyP pool might buffer the cytosol against sudden drops in P_i_ and delay the onset of the P_i_ starvation response (6, 14). In line with this, our metabolomic analysis revealed exaggerated metabolic changes when the polyP-deficient vtc4Δ mutant was starved for P_i_. This supports a significant role of acidocalcisome-like vacuoles in P_i_ homeostasis, which may provide the proposed buffer for cytosolic P_i_. Such a buffer is of obvious relevance for an ordered transition into P_i_-starvation and cell cycle arrest (44, 87). Surprisingly, metabolic features of P_i_ starvation were observed in the vtc4Δ mutant already under high P_i_ conditions. This phenotype may again reflect the buffering function of polyP. This function may become important even on high P_i_ media because the duplication of all nucleic acids and phospholipids generates a very high need for P_i_ in S-phase. It has been proposed that this need can transiently exceed the maximal import capacity of cells, necessitating the vacuolar polyP store to cover the deficit (43, 44). An explanation of the starvation features of vtc4Δ from this perspective is, however, only partially satisfactory for our dataset because vtc4Δ on high P_i_ shows numerous but not all features associated with P_i_ starvation. The trend for some metabolites was even inversed, such as for adenosine, guanosine, orotic acid, acetyl-CoA and phosphoenolpyruvate. This suggests that polyP may have additional metabolic functions that go beyond those of a P_i_ buffer for the cytosol. There is potential for this because polyP has a significant role for the storage of cations, such as for Zn^2+^, Ca^2+^, Mn^2+^ and Mg^2+^ (88–91), but also for cation uptake, as shown for Mg^2+^ (92). Furthermore, polyP may affect cellular signaling, influencing the stress response (93–96). Here, it may even have direct impact, such as through polyphosphorylation of lysine residues, which modifies yeast topoisomerase 1 (Tpo1) and nuclear signal recognition factor 1 (Nsr1) (97, 98).

In sum, our observations favor a model where lack of P_i_ induces differential metabolic changes, which together promote the P_i_ starvation response. Beginning P_i_ scarcity could be diagnosed through moderate declines in ATP and inositol pyrophosphates, leading the cells to partially activate P_i_ scavenging systems and maintain normal growth. Profound P_i_ starvation entails numerous additional metabolic changes, such as through AICAR, SAM, and strong reductions in ATP and inositol pyrophosphates. These changes may fully stimulate the transcriptional P_i_ starvation and stress responses in a combinatorial manner. We consider this latter point as an attractive hypothesis addressing the problem of how cells measure P_i_. Given that P_i_ is present in the cytosol in millimolar concentrations it is can hardly be envisioned that it be “measured” by specific binding to a low affinity receptor, which might be susceptible to competition by numerous other compounds. Coincidence detection through a network of P_i_-dependent metabolic reactions could, however, generate such specificity, even for a highly abundant ligand such as P_i_. Therefore, we favor this model of P_i_ detection.

## Materials and methods

### Yeast strains and media

The *S. cerevisiae* strains used in this study are listed in Table 4. Endogenous GFP tagging was performed as described previously (99). The yEGFP-CaURA3 was PCR amplified from the plasmid pKT209 by introducing 40 bp homology before and after the stop codon of the Pho4 gene. Gene deletion was conducted based on the CRISPR-Cas9 system as described previously (100). The single-guide RNA (sgRNA) was cloned into the sgRNA expression vector and co-transformed into yeast cells with hybridized double-stranded oligonucleotides, which contain 40 bp homology of each side before the start codon and after the stop codon of Vtc4 gene as templates for homologous recombination. After transformation, positive colonies were selected by colony PCR and sequencing. PCR primers used for genetic manipulation are listed in Table 5.

**Table 4.** Yeast strains used in this study.

| Strain | Genotype | Source |
| --- | --- | --- |
| BY4741 | *MATa; his3Δ1; leu2Δ0; met15Δ0; ura3Δ0* | Euroscarf |
| BY4741 Pho4-yeGFP | *MATa; his3Δ1; leu2Δ0; met15Δ0; ura3Δ0; Pho4-yeGFP::CaURA* | This study |
| BY4741 *vtc4Δ* | *MATa; his3Δ1; leu2Δ0; met15Δ0; ura3Δ0; vtc4Δ* | This study |

**Table 5.**
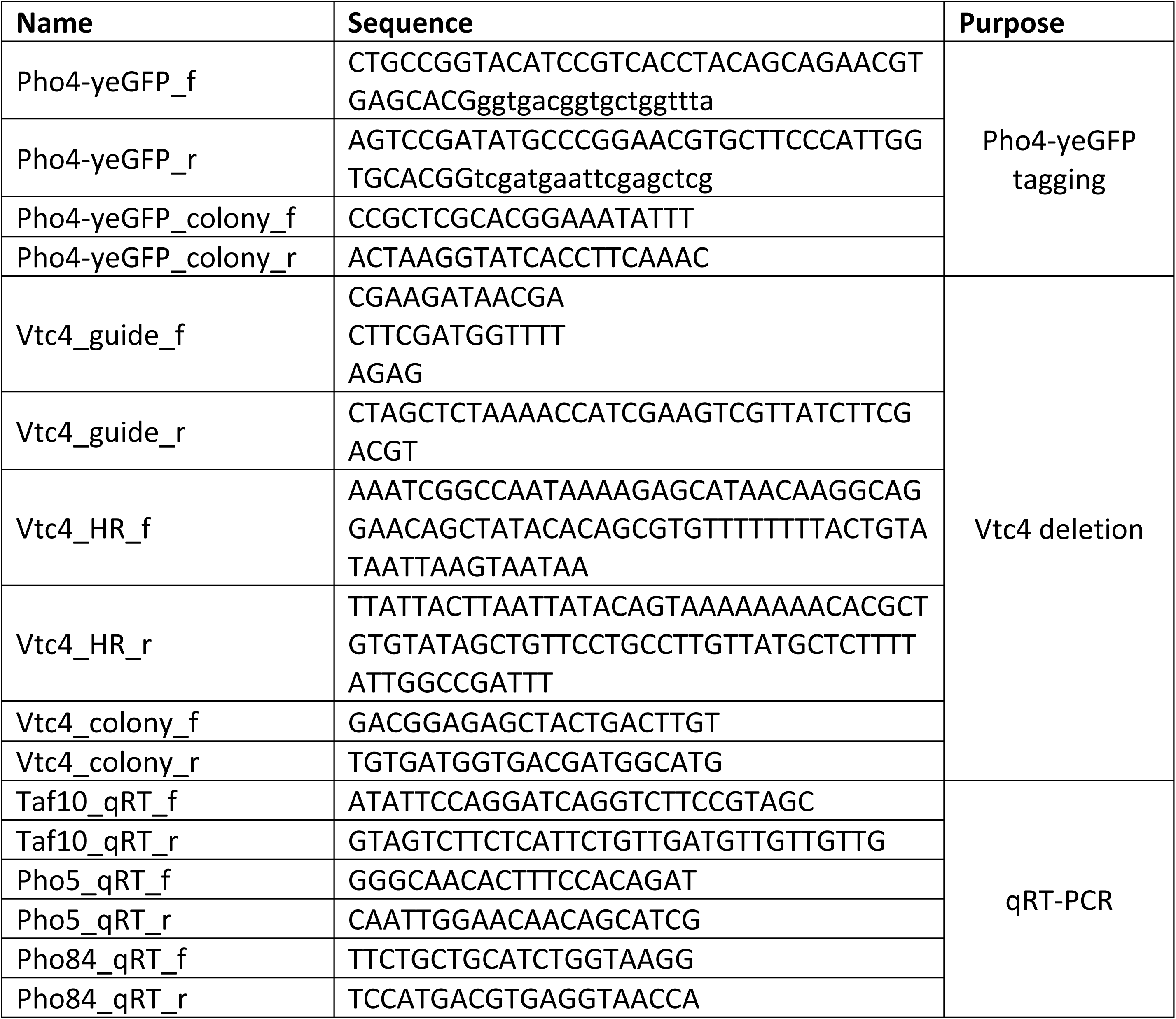
Primers used in this study.

Synthetic complete (SC) medium was prepared using yeast nitrogen base without phosphate (Formedium, UK). Desired phosphate concentration was adjusted by adding KH_2_PO_4_. The potassium concentration was controlled by adding KCl instead of KH_2_PO_4_.

### Growth assay

Yeast cells were grown overnight in P_i_-rich SC medium 10 mM P_i_ until the logarithmic growth phase (OD_600nm_ ≈ 1). Cells were divided into 6 tubes, centrifuged, and washed by SC medium containing different concentrations of P_i_ (10 mM, 1 mM, 0.5 mM, 0.25 mM, 0.125 mM, and 0 mM). After two rounds of washing, cells were inoculated to relevant SC media (OD600_nm_ ≈ 0.05) and grown at 30 °C in a shaking incubator. To assess the growth, OD_600nm_ was monitored at different time points (0 h, 0.5 h, 1 h, 2 h, 4 h, and 24 h).

### Acid phosphatase assay

Acid phosphatase assay was performed as described (101). Cells were grown in the same manner for growth assay as described above. At each time point, 0.2 OD_600nm_ units of cells were harvested by centrifugation and resuspended in 250 µl of 0.1 M sodium acetate (pH 4.2) and 250 µl of freshly prepared 9 mg/ml p-nitrophenyl phosphate. The mixture was incubated at 37 °C for 9 min and 800 µl of 1.4 M Na_2_CO_3_ was added to stop the reaction. After centrifugation, OD_420nm_ was measured from the supernatant as acid phosphatase activity.

### Fluorescence microscopy

Cells in the logarithmic phase were inoculated into SC media containing 10 mM, 0.5 mM, or 0 mM P_i_ and incubated for 8 h at 30 °C. grown in the same manner for growth assay as described above. Fluorescence images were obtained by a Nikon Eclipse Ti2/Yokogawa CSU-X1 Spinning Disk microscope with two Prime BSI sCMOS cameras (Teledyne Photometrics, USA), a LightHub Ultra ® Laser Light (Omicron Laserage, Germany), and an Apo TIRF 100x/1.49 Oil lens (Nikon, Japan). Experiments were repeated and representative images were shown in the manuscript.

### RNA extraction and qRT-PCR

Total RNA was extracted from 10 OD_600nm_ units of yeast cells by RNeasy kits (Qiagen, Germany) according to the manufacturer’s instructions. 1 µg of total RNA was used for cDNA synthesis using RevertAid reverse transcriptase (Thermo Fisher Scientific, USA). Gene expression level was quantitatively monitored using real-time PCR (LightCycler 480, Roche, Switzerland) with SYBR Green I Master (Roche, Switzerland). Gene expression was normalized by TATA-binding protein-Associated Factor Taf10 transcript as an internal control. Primers used for qRT-PCR are listed in Table 5. The mean and standard deviation of gene expression was calculated from three biological replicates with three technical replicates.

### PolyP measurement

PolyP levels were evaluated from cells using the direct 4’-6-diamidino-2-phenylindole (DAPI) assay [62]. Cells were grown in P_i_-rich SC medium until the logarithmic phase and inoculated to SC media containing different concentrations of P_i_. After 8 h of incubation, 0.5 OD_600nm_ units of cells were harvested and washed with 50 mM HEPES-KOH (pH 7.5). The cell pellet was resuspended in 400 µl of DAPI buffer containing 20 mM HEPES-KOH (pH 6.8), 150 mM KCl, and 10 µM DAPI. After two rounds of freeze-thaw in liquid N_2_, samples were centrifuged for 2 min at 13,000 rpm. The supernatant was diluted with DAPI buffer (1:20) and 200 µl of diluted samples was transferred to black 96-well polypropylene plates. The DAPI fluorescence was measured by SpectraMax Gemini EM (Molecular Devices, USA) with excitation and emission of 420 nm and 550 nm respectively (102, 103).

### Extraction of inositol pyrophosphates for quantification

To extract InsPPs, 1 ml of cell culture grown in the SC medium containing different P_i_ levels (10 mM, 0.5 mM, and 0 mM) for 8 h was transferred into a microcentrifuge tube and mixed with 100 µl of 11 M perchloric acid. The mixture was snap-frozen in liquid N_2_ and centrifuged (3 min, 13,000 rpm, 4 °C) and then the supernatant was transferred into a new tube. Titanium dioxide (TiO_2_) beads (GL Sciences, Japan) were added after two rounds of washing with H_2_O and 1 M perchloric acid (1.5 mg of beads/sample). The sample with TiO_2_ beads was gently rotated for 15 min at 4 °C and centrifuged (1 min, 13,000 rpm, 4 °C). The beads were washed twice using 500 µl of 1 M perchloric acid and resuspended by 300 µl of 3 % (v/v) NH_4_OH followed by gentle rotation at room temperature. After centrifugation (1 min, 13,000 rpm), the eluents were transferred into a new tube and completely dried under the SpeedVac (Labogene) at 42 °C.

### Analysis of PP-InsPs

The amount of InsPPs was quantified by capillary electrophoresis coupled to electrospray ionization mass spectrometry (CE-ESI-MS) as described previously (104–106). E-ESI-MS analysis was performed on an Agilent 7100 CE coupled to triple quadrupole mass spectrometer (QqQ MS) Agilent 6495c, equipped with an Agilent Jet Stream (AJS) electrospray ionization (ESI) source. Stable CE-ESI-MS coupling was enabled by commercial sheath liquid coaxial interface, with an isocratic LC pump constantly delivering the sheath-liquid.

All experiments were performed with a bare fused silica capillary with a length of 100 cm and 50 μm internal diameter. 35 mM ammonium acetate titrated by ammonia solution to pH 9.7 was used as background electrolyte (BGE). The sheath liquid was composed of a water-isopropanol (1:1) mixture, with a flow rate of 10 µL/min. 15 µL InsPs extracts was spiked with 0.75 µL isotopic internal standards mixture (2 µM [^13^C_6_]1,5-InsP_8_, 10 µM [^13^C_6_]5-InsP_7_, 10 µM [^13^C_6_]1-InsP_7_ and 40 µM [^13^C_6_]InsP_6_). Samples were introduced by applying 100 mbar pressure for 10 s (20 nL). A separation voltage of +30 kV was applied over the capillary, generating a constant CE current at around 19 µA.

The MS source parameters setting were as follows: nebulizer pressure was 8 psi, sheath gas temperature was 175 °C and with a flow at 8 L/min, gas temperature was 150 °C and, with a flow of 11 L/min, the capillary voltage was −2000 V with nozzle voltage 2000V. Negative high pressure RF and low pressure RF (Ion Funnel parameters) was 70V and 40V, respectively. Parameters for MRM transitions are as follows:

| Compound | Precursor Ion | Product Ion | dwell | CE (V) | Cell Acc (V) | Polarity |
| --- | --- | --- | --- | --- | --- | --- |
| [$^{13}\text{C}_6$]InsP<sub>8</sub> | 411.9 | 362.9 | 80 | 10 | 1 | Negative |
| InsP<sub>8</sub> | 408.9 | 359.9 | 80 | 10 | 1 | Negative |
| [$^{13}\text{C}_6$]InsP<sub>7</sub> | 371.9 | 322.9 | 80 | 10 | 1 | Negative |
| InsP<sub>7</sub> | 368.9 | 319.9 | 80 | 10 | 1 | Negative |
| [$^{13}\text{C}_6$]InsP<sub>6</sub> | 331.9 | 486.9 | 80 | 17 | 1 | Negative |
| InsP<sub>6</sub> | 328.9 | 480.9 | 80 | 17 | 1 | Negative |

### Preparation of yeast metabolite extracts

0.5 OD_600_ units of log-phase yeast cells grown in synthetic complete (SC) media were collected by the vacuum-filtration method using a polytetrafluoroethylene membrane filter (1.2 μm; Piper Filter GmbH, Germany) (52). Yeast cells on PTFE membrane were resuspended with MeOH:H_2_O (4:1, v/v) mixture. The samples were homogenized by the Cryolys Precellys 24 sample homogenizer (Bertin Technologies, USA) with ceramic beads. Homogenized extracts were centrifuged for 15 min at 4000 g at 4°C. The precipitated protein pellets were used to measure total protein concentration and the supernatant was collected and evaporated in a vacuum concentrator (LabConco, USA). Dried sample extracts were resuspended in MeOH:H_2_O (4:1, v/v) mixture based on the total protein content.

### LC-MS

Untargeted metabolite profiling was performed by hydrophilic interaction liquid chromatography coupled to tandem mass spectrometry (HILIC – MS/MS) in both positive and negative ionization modes using a 6495 triple quadrupole system (QqQ) interfaced with 1290 UHPLC system (Agilent Technologies, USA) [64]. In positive mode, the chromatographic separation was carried out in an Acquity BEH Amide, 1.7 μm, 100 mm x 2.1 mm I.D. column (Waters, USA). The mobile phase was composed of A = 20 mM ammonium formate and 0.1 % formic acid in water and B = 0.1 % formic acid in acetonitrile. The linear gradient elution from 95% B (0-1.5 min) down to 45% B was applied (1.5-17 min) and these conditions were held for 2 min. Then initial chromatographic conditions were maintained as a post-run during 5 min for column re-equilibration. The flow rate was 400 μl/min, column temperature 25°C, and sample injection volume 2 μl. Electro-spray ionization (ESI) source conditions were set as follows: dry gas temperature 290°C, nebulizer 35 psi and flow 14 l/min, sheath gas temperature 350°C and flow 12 l/min, nozzle voltage 0 V, and capillary voltage 2000 V. Dynamic multiple reaction monitoring (DMRM) was used as acquisition mode with a total cycle time of 600 ms. Optimized collision energies for each metabolite were applied. In negative mode, a SeQuant ZIC-pHILIC (100 mm, 2.1 mm I.D. and 5 μm particle size, Merck, Germany) column was used. The mobile phase was composed of A = 20 mM ammonium acetate and 20 mM ammonium hydroxide in water at pH 9.7 and B = 100% acetonitrile. The linear gradient elution from 90% (0-1.5 min) to 50% B (8-11 min) down to 45% B (12-15 min). Finally, the initial chromatographic conditions were established as a post-run during 9 min for column re-equilibration. The flow rate was 300 μl/min, column temperature 30°C and sample injection volume 2 μl. ESI source conditions were set as follows: dry gas temperature 290°C and flow 14 l/min, sheath gas temperature 350°C, nebulizer 45 psi, and flow 12 l/min, nozzle voltage 0 V, and capillary voltage −2000V. DMRM was used as an acquisition mode with a total cycle time of 600 ms. Optimized collision energies for each metabolite were applied.

### Data pre-processing

Raw LC-MS/MS data were processed using the Agilent Quantitative analysis software (version B.07.00 MassHunter, Agilent Technologies, USA). Relative quantification of metabolites was based on Extracted Ion Chromatogram (EIC) areas for the monitored MRM transitions. The obtained results were exported to “R” software (http://cran.r-project.org/) and signal intensity drift correction was done within the LOWESS/Spline normalization program followed by noise filtering (CV(QC features) > 30%).

### Statistical analysis of metabolite profiling

Statistical analyses of metabolomic data were performed by MetaboAnalyst 5.0 (https://www.metaboanalyst.ca/) [65]. Before analysis, signal intensity data were median-normalized, log-transformed, and mean-centered using the auto-scaling method. PLS-DA of the first metabolic profiling with different P_i_ conditions was conducted by considering the P_i_ concentration order. The heatmap of the correlation matrix between metabolites of different P_i_ conditions was calculated by the Pearson r correlation coefficient. Volcano plot analysis was performed by a two-sample t-test. The metabolites showing p < 0.1 with |FC| > 1.5 (10 mM P_i_ vs 0.5 mM P_i_) or |FC| > 2 (10 mM P_i_ vs 0 mM P_i_) were considered as statistically meaningful metabolites. Results of volcano plot analysis were exported and visualized with GraphPad Prism 9 (GraphPad Software, USA). For the second metabolic profiling with wildtype and vtc4Δ comparison, the prominent outliers were removed after PLS-DA for further analyses. A two-way ANOVA followed by false discovery rate correction (p < 0.05) was performed to investigate metabolite variabilities between two different factors, genotype (wildtype and vtc4Δ) and P_i_ conditions (10 mM P_i_ and 0 mM P_i_), and their interaction. A hierarchical clustering heatmap was generated using Euclidean distance measure with Ward’s clustering method. The top 50 significant metabolites from ANOVA were selected for visualization.

### Pathway analysis and metabolite set enrichment analysis

Pathway analysis performed using MetaboAnalyst 5.0 based on the metabolites that statistically significantly increased or decreased in 0.5 mM P_i_ (|FC| > 1.5; p < 0.1) or 0 mM P_i_ (|FC| > 2; p < 0.1) conditions compared to 10 mM P_i_ conditions. Hypergeometric test and relative-betweenness centrality were used for the enrichment method and topology analysis, respectively. Metabolite set enrichment analysis was performed by MetaboAnalyst 5.0 based on 84 metabolite sets of Kyoto Encyclopedia of Genes and Genomes (KEGG) human metabolic pathways. Results of pathway analysis and metabolite set enrichment analysis were exported and visualized with GraphPad Prism 9.

## Acknowledgements

We thank the metabolomics facility of UNIL for support with the metabolite analyses. This project was funded through grants from the ERC (No. 788442 to AM) and HFSP (LT000588/2019 to GK).

---

**Supplementary Figure 1.**
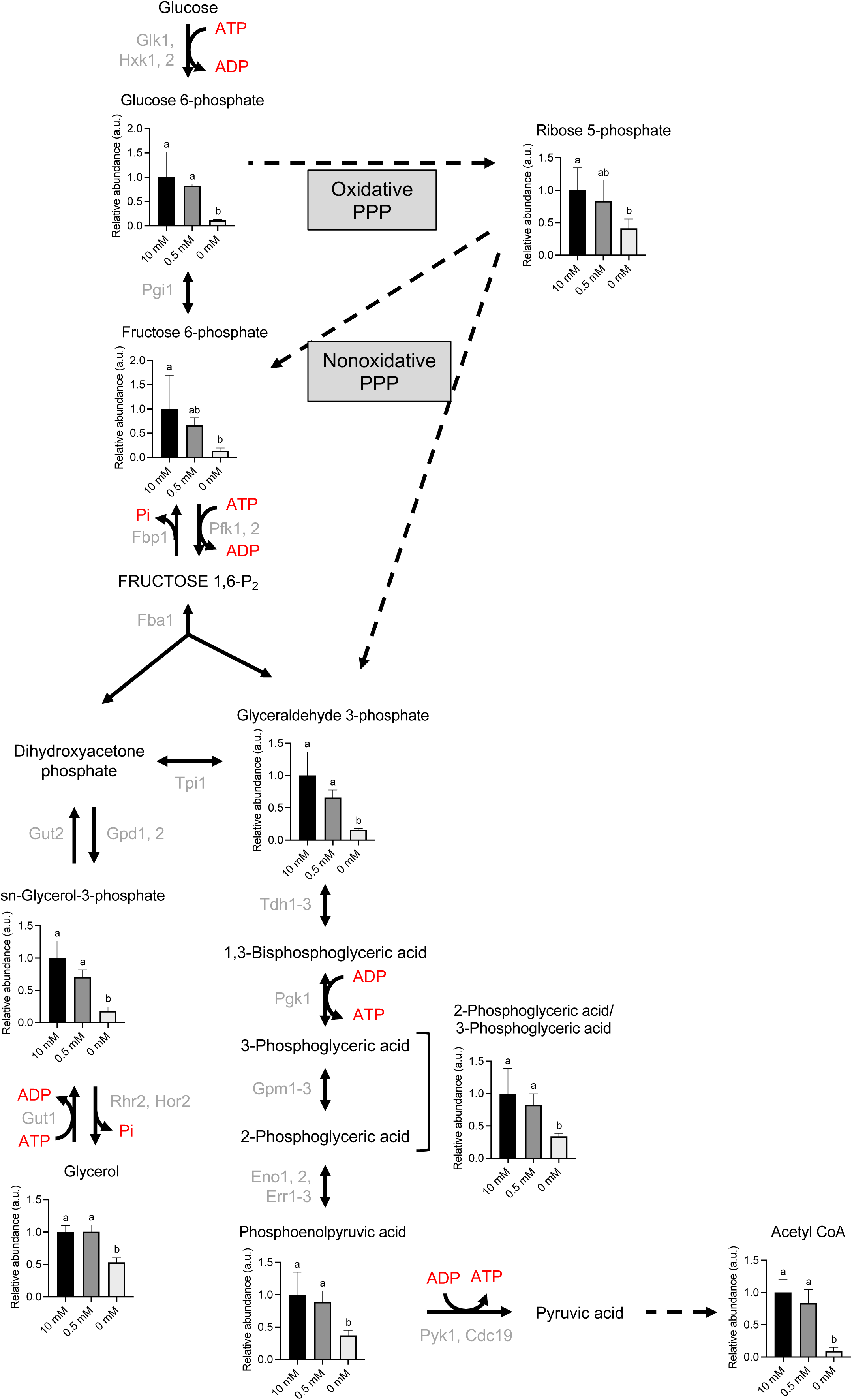
Metabolic changes in glycolysis under different Pi conditions. Metabolomic data of yeast under different Pi conditions were presented on a metabolic pathway. Pi-containing substrates were marked as red. Grey indicates enzymes involved in the reaction. Different letters on the graph indicate a significance difference by Tukey-Kramer test (p < 0.05).

**Supplementary Figure 2.**
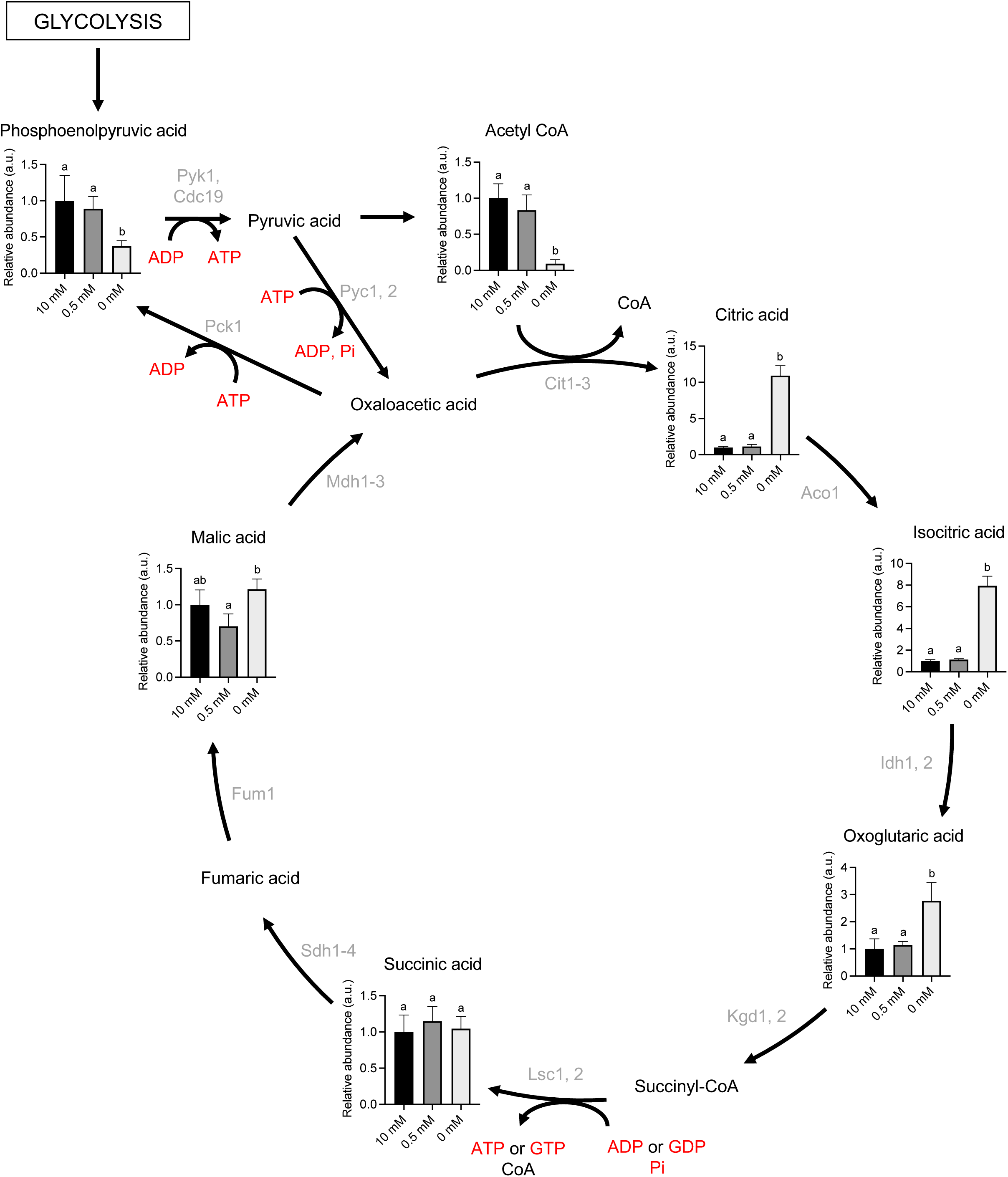
Metabolic changes in TCA cycle under different Pi conditions. Metabolomic data of yeast under different Pi conditions were presented on a metabolic pathway. Pi-containing substrates were marked as red. Grey indicates enzymes involved in the reaction. Different letters on the graph indicate a significance difference by Tukey-Kramer test (p < 0.05).

**Supplementary Figure 3.**
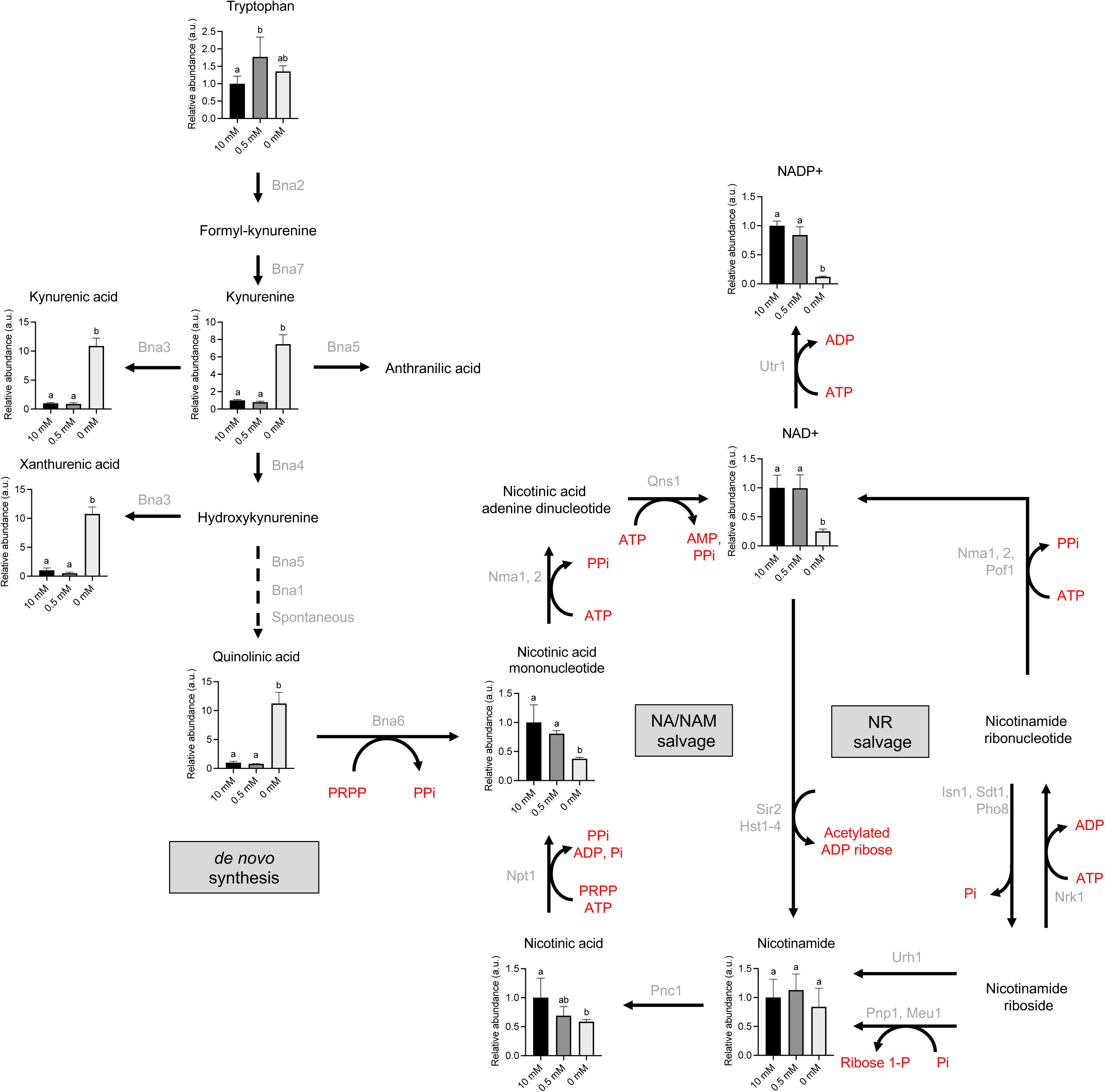
Metabolic changes in nicotinate and nicotinamide pathway under different Pi conditions. Metabolomic data of yeast under different Pi conditions were presented on a metabolic pathway. Pi-containing substrates were marked as red. Grey indicates enzymes involved in the reaction. Different letters on the graph indicate a significance difference by Tukey-Kramer test (p < 0.05).

**Supplementary Figure 4.**
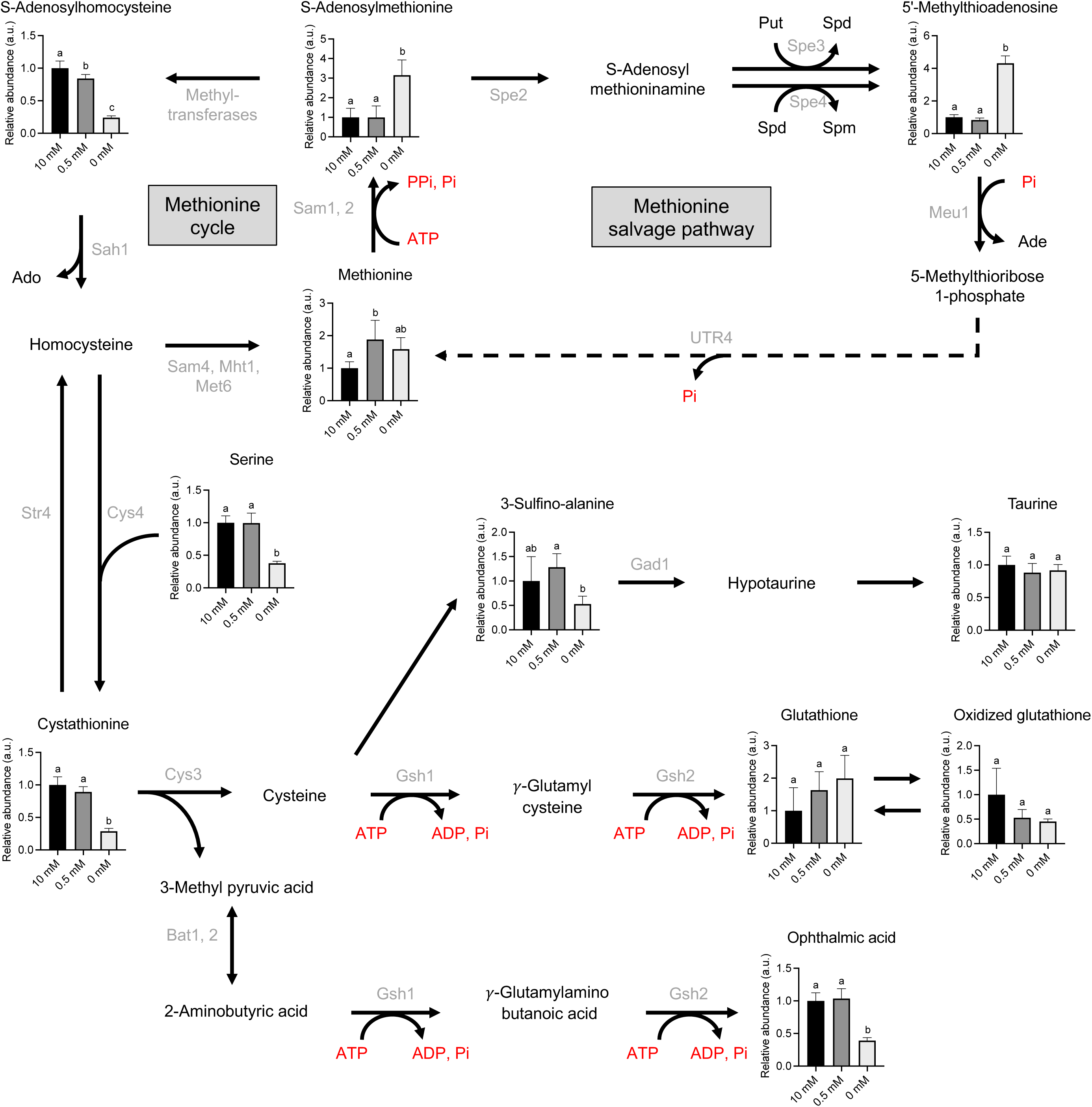
Metabolic changes in methionine and cysteine metabolism pathway under different Pi conditions. Metabolomic data of yeast under different Pi conditions were presented on a metabolic pathway. Pi-containing substrates were marked as red. Grey indicates enzymes involved in the reaction. Different letters on the graph indicate a significance difference by Tukey-Kramer test (p < 0.05).

**Supplementary Figure 5.**
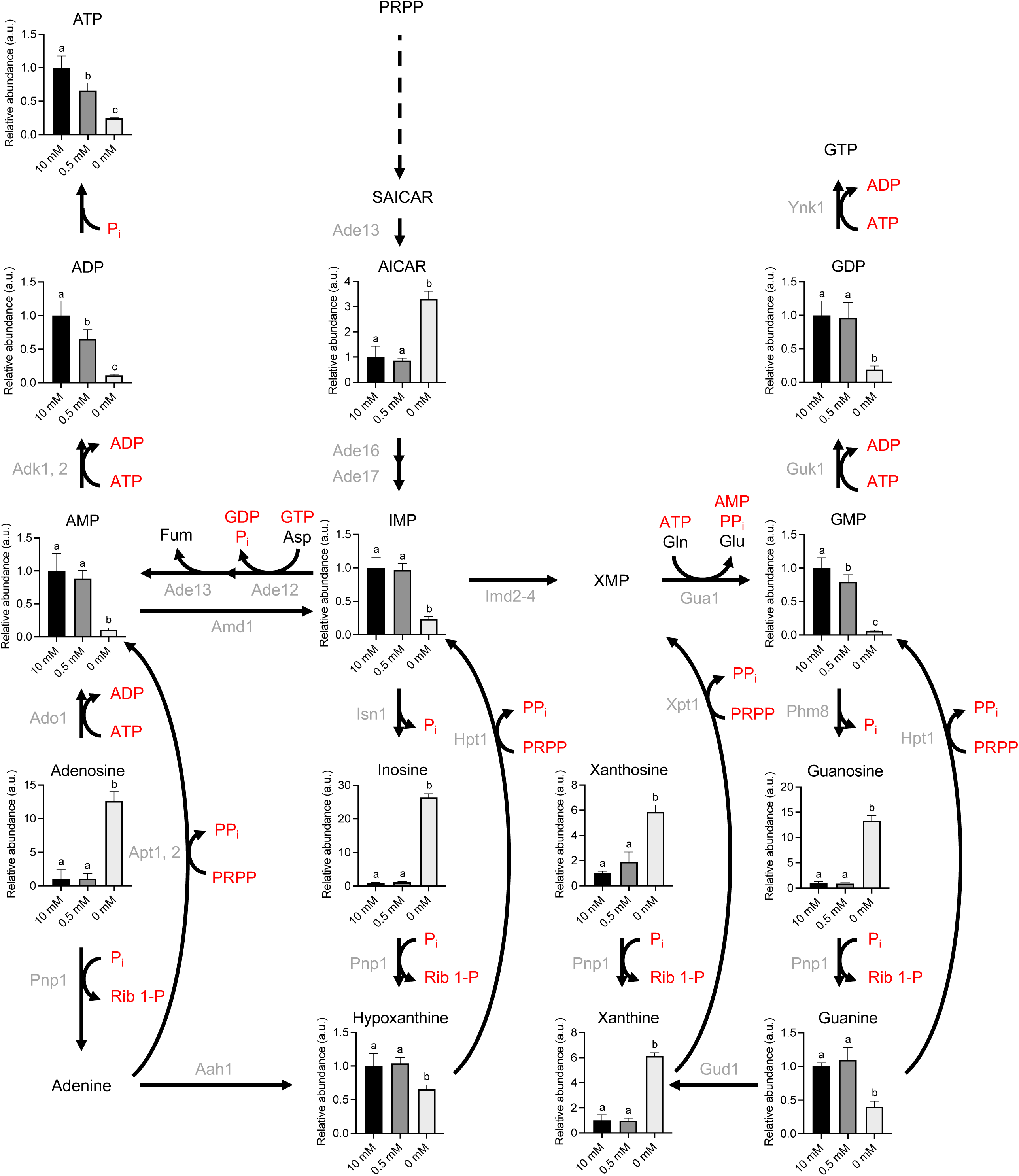
Metabolic changes in purine metabolism pathway under different P_i_ conditions. Metabolomic data of yeast under different Pi conditions were presented on a metabolic pathway. Pi-containing substrates were marked as red. Grey indicates enzymes involved in the reaction. Different letters on the graph indicate a significance difference by Tukey-Kramer test (p < 0.05).

**Supplementary Figure 6.**
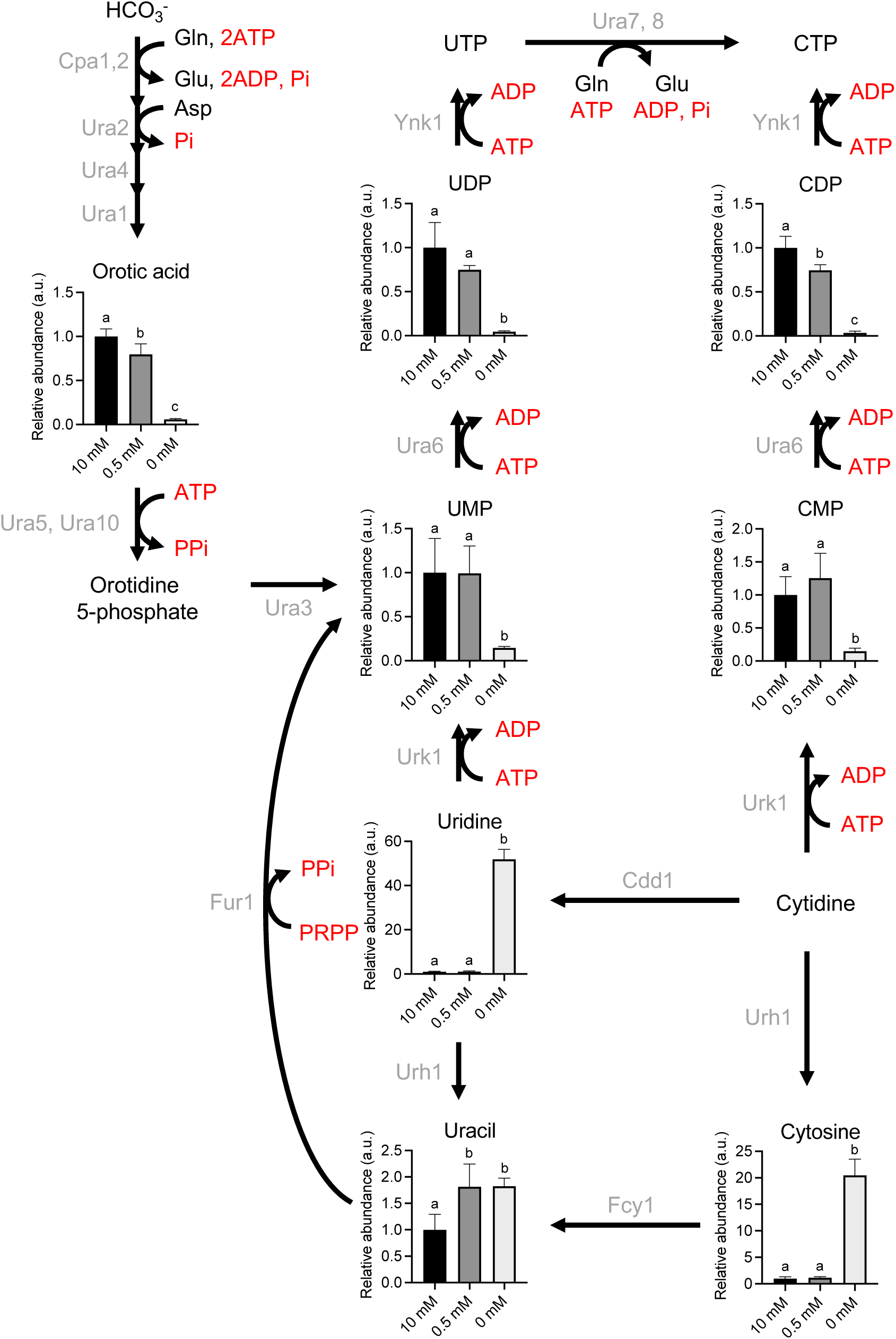
Metabolic changes in pyrimidine metabolism pathway under different Pi conditions. Metabolomic data of yeast under different Pi conditions were presented on a metabolic pathway. Pi-containing substrates were marked as red. Grey indicates enzymes involved in the reaction. Different letters on the graph indicate a significance difference by Tukey-Kramer test (p < 0.05).

**Supplementary Figure 7.**
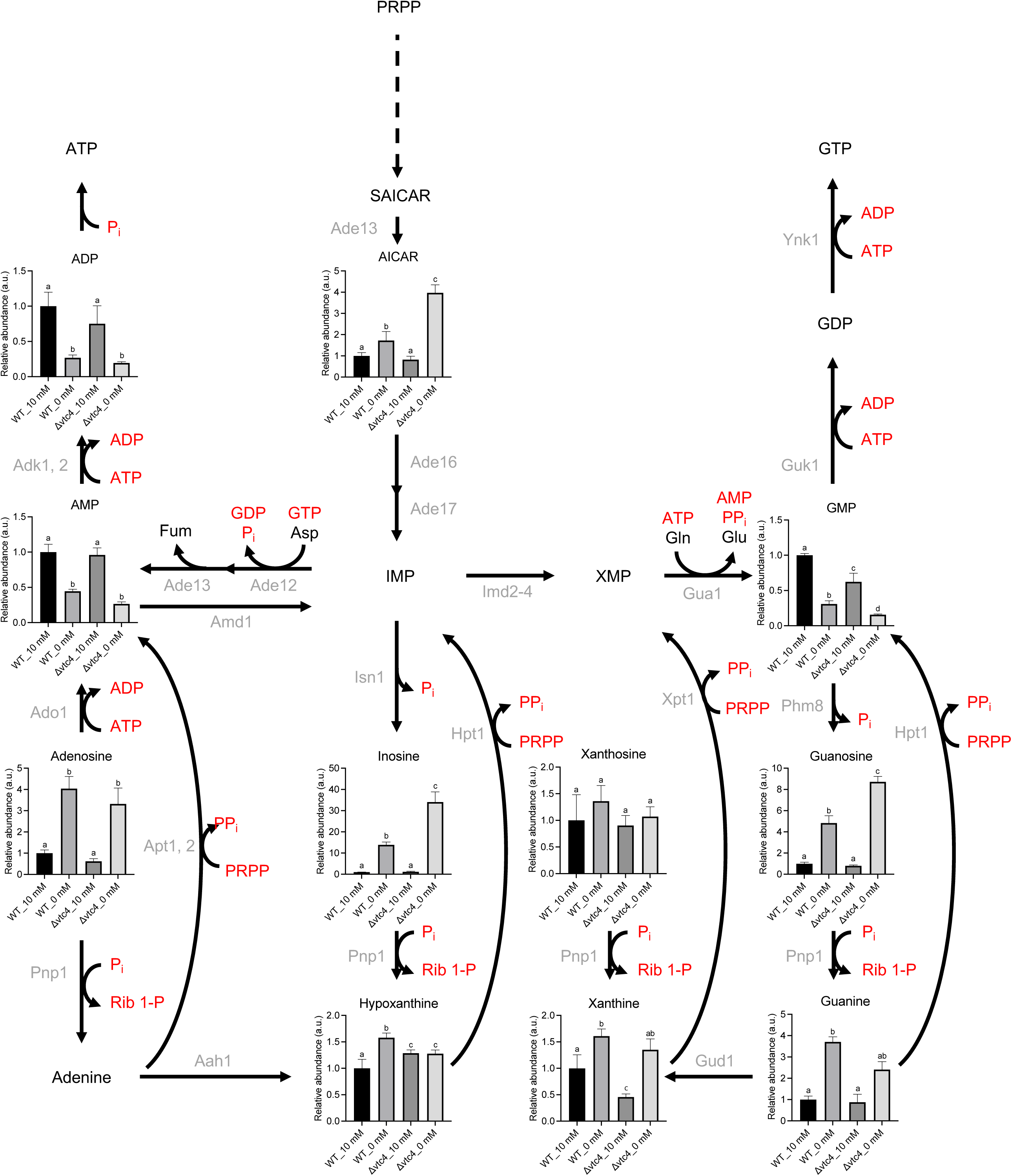
Metabolic changes in purine metabolism pathway under Pi starvation in wild type and vtc4Δ. Metabolomic data of yeast under Pi starvation in wild type and vtc4Δ were presented on a metabolic pathway. Pi-containing substrates were marked as red. Grey indicates enzymes involved in the reaction. Different letters on the graph indicate a significance difference by Tukey-Kramer test (p < 0.05).

**Supplementary Figure 8.**
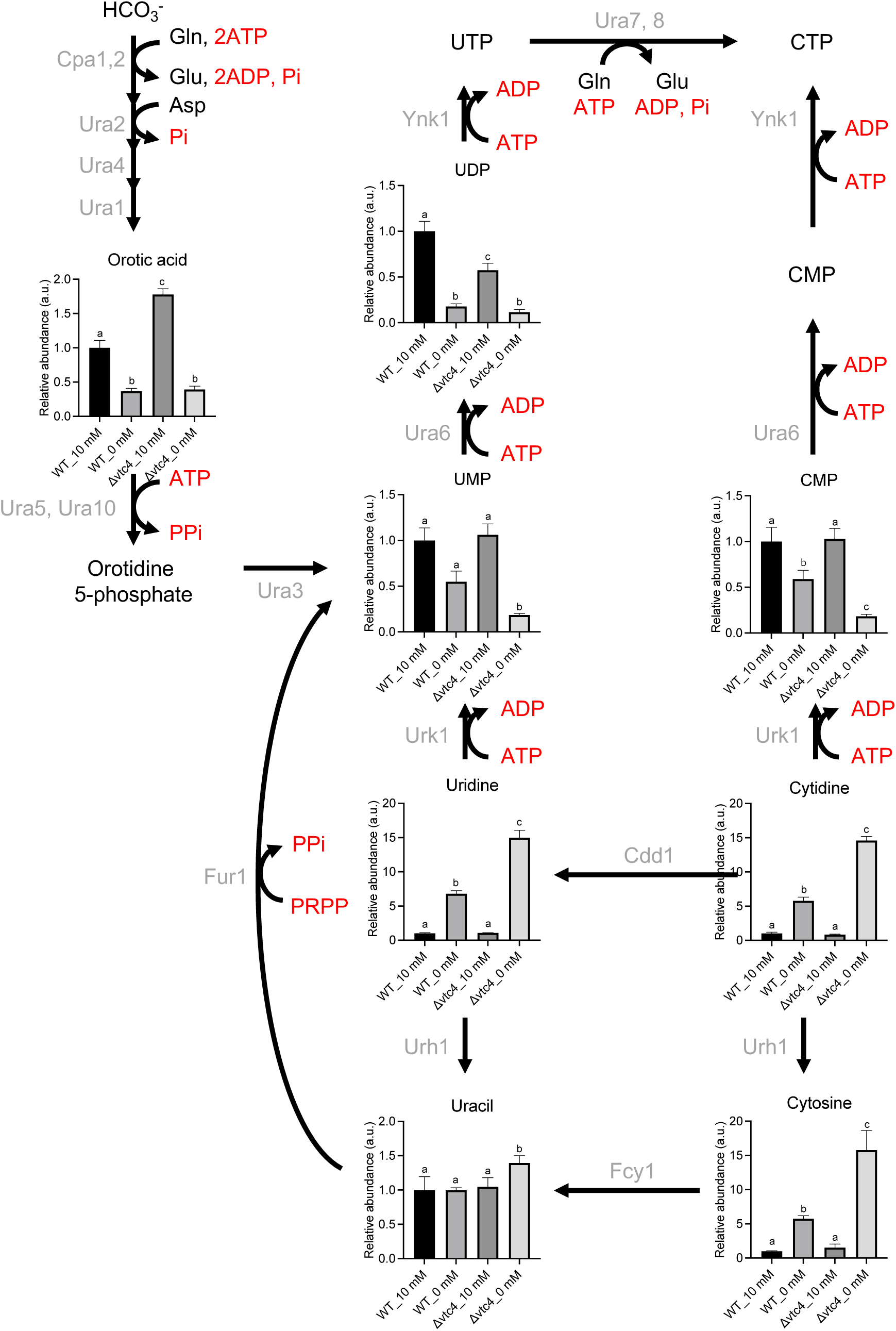
Metabolic changes in pyrimidine metabolism pathway under Pi starvation in wild type and vtc4Δ. Metabolomic data of yeast under Pi starvation in wild type and vtc4Δ were presented on a metabolic pathway. Pi-containing substrates were marked as red. Grey indicates enzymes involved in the reaction. Different letters on the graph indicate a significance difference by Tukey-Kramer test (p < 0.05)

**Supplementary Figure 9.**
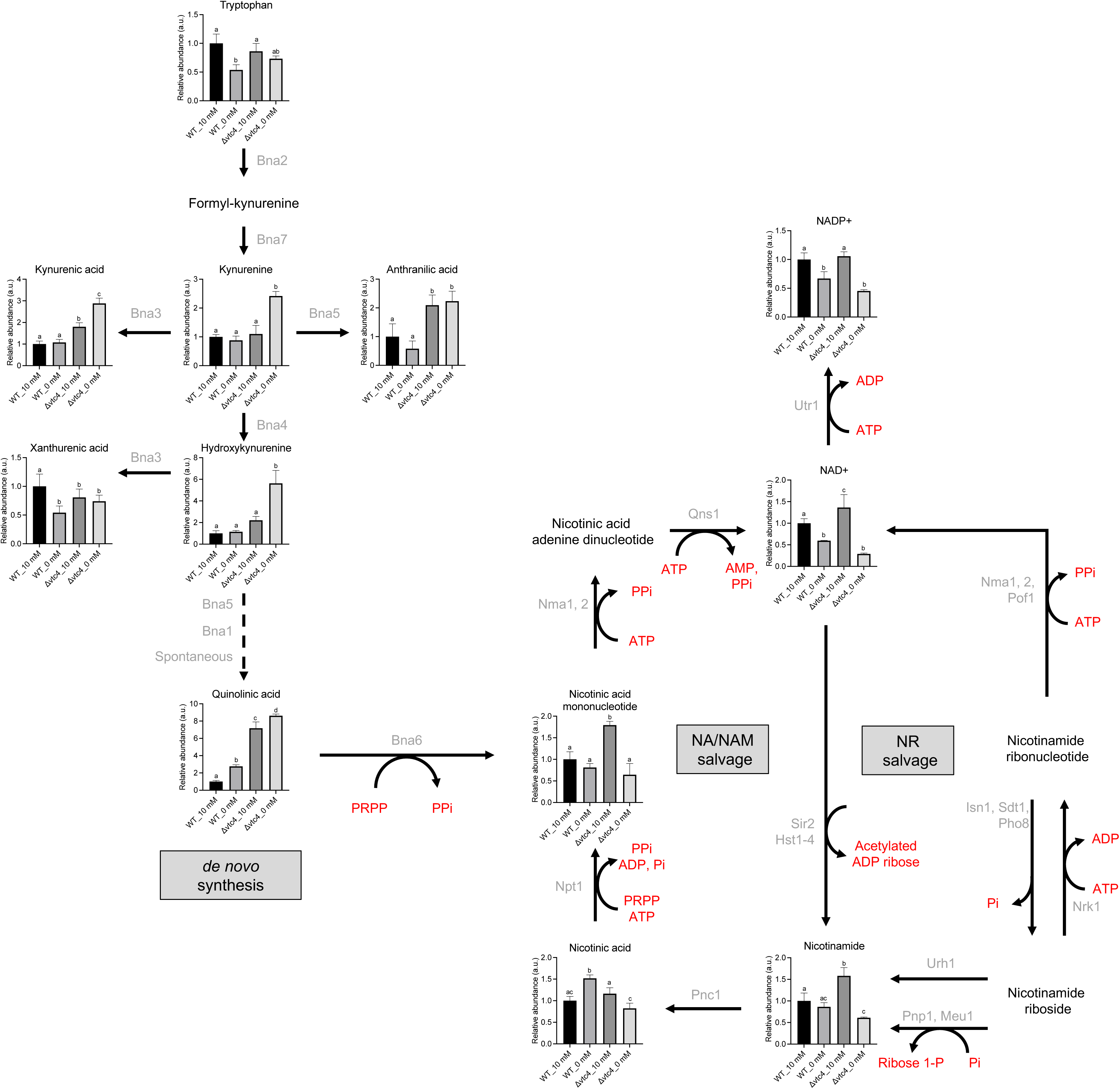
Metabolic changes in nicotinate and nicotinamide pathway under Pi starvation in wild type and vtc4Δ. Metabolomic data of yeast under Pi starvation in wild type and vtc4Δ were presented on a metabolic pathway. Pi-containing substrates were marked as red. Grey indicates enzymes involved in the reaction. Different letters on the graph indicate a significance difference by Tukey-Kramer test (p < 0.05).

**Supplementary Figure 10.**
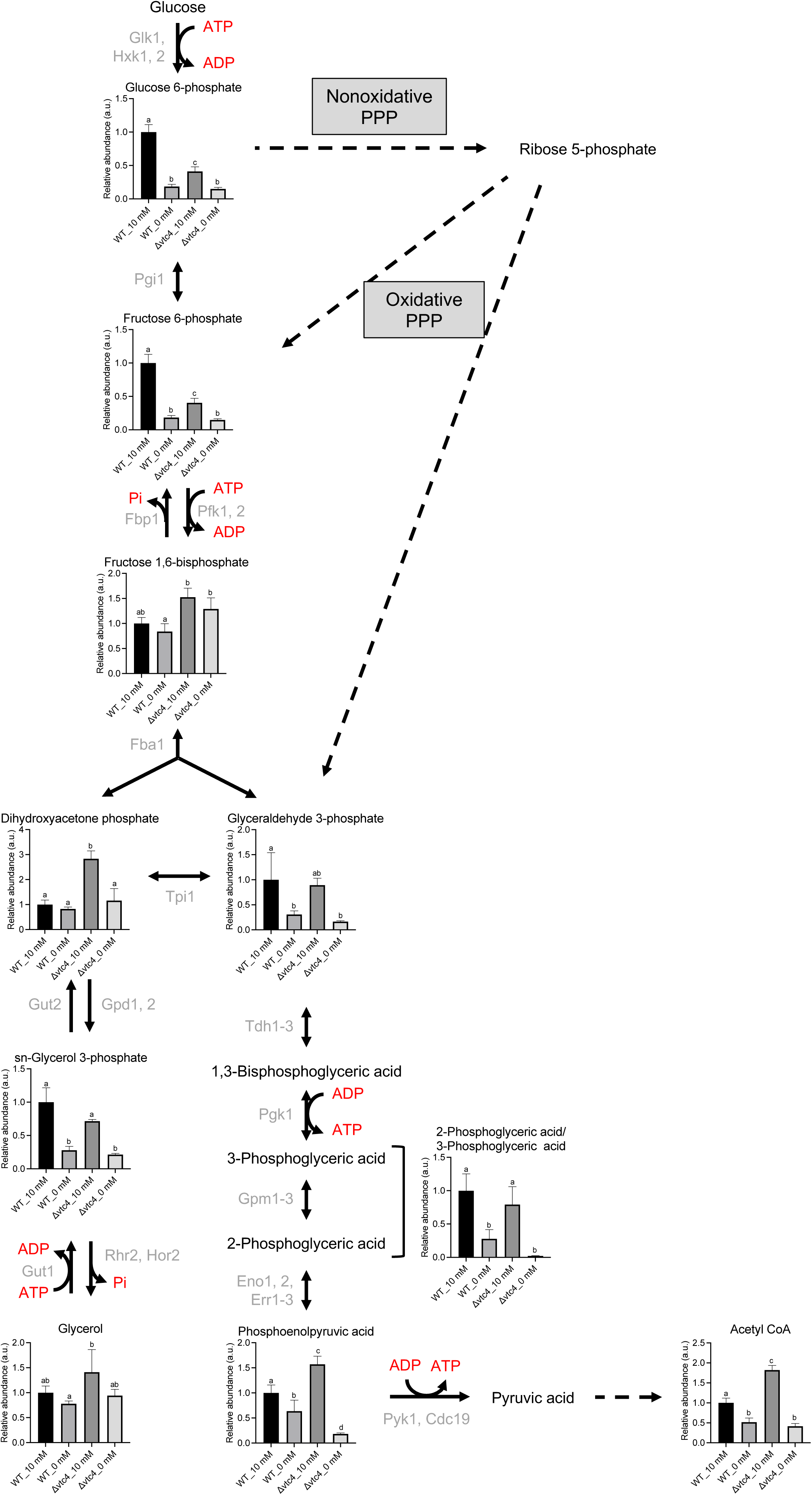
Metabolic changes in glycolysis under Pi starvation in wild type and vtc4Δ. Metabolomic data of yeast under Pi starvation in wild type and vtc4Δ were presented on a metabolic pathway. Pi-containing substrates were marked as red. Grey indicates enzymes involved in the reaction. Different letters on the graph indicate a significance difference by Tukey-Kramer test (p < 0.05).

